# Novel porcine model reveals two distinct LGR5 cell types during lung development and homeostasis

**DOI:** 10.1101/2022.12.09.516617

**Authors:** Kathryn M. Polkoff, Ross Lampe, Nithin K. Gupta, Yanet Murphy, Jaewook Chung, Amber Carter, Jeremy M. Simon, Katherine Gleason, Adele Moatti, Preetish K. Murthy, Laura Edwards, Alon Greenbaum, Aleksandra Tata, Purushothama Rao Tata, Jorge A. Piedrahita

## Abstract

Cells expressing LGR5 play a pivotal role in homeostasis, repair, and regeneration in multiple organs including skin and gastrointestinal tract, yet little is known about their role in the lung. Findings from mice, a widely used animal model, suggest that lung LGR5 expression differs from that of humans. In this work, using a new transgenic pig model, we identify two main populations of LGR5^+^ cells in the lung that are conserved in human, but not mouse lungs. Using RNA sequencing, 3D imaging and organoid models, we determine that in the fetal lung, epithelial LGR5 expression is transient in a subpopulation of SOX9^+^/ETV^+^/SFTPC^+^ progenitor lung tip cells. In contrast, epithelial LGR5 expression is absent from postnatal lung, but is reactivated in bronchioalveolar organoids derived from basal airway cells. We also describe a separate population of mesenchymal LGR5^+^ cells that surrounds developing and mature airways, lies adjacent to airway basal cells, and is closely associated with nerve fibers. Transcriptionally, mesenchymal LGR5^+^ cells include a subset of peribronchial fibroblasts (PBF) that express unique patterns of *SHH, FGF, WNT* and *TGF-*β signaling pathway genes. These results support distinct roles for LGR5^+^ cells in the lung and describe a physiologically relevant animal model for further studies on the function of these cells in repair and regeneration.

## Introduction

LGR5 is a known potentiator of WNT signaling (1) is often involved in developmental processes and regeneration, and marks progenitor populations in several organs including GI and skin (2,3). In humans, study of LGR5^+^ cells have been sparse due to lack of reliable antibodies. WNT signaling, in addition, is vital to lung development (4,5) and repair and regeneration (6). And RSPO2, a ligand for LGR5, was has a vital role in lung branching morphogenesis, and the limited lung growth resulting from inactivation of *RSPO2* was correlated with reduced canonical WNT signaling (7). These observations suggest a potential role for LGR5 signaling in lung development. While distribution of lung Lgr5^+^ cells have not been described in fetal mice, in humans, using scRNA-seq, an *LGR5^+^* epithelial population was reported in the elongating bud tip of the fetal lung (8). In the postnatal lung, a study of *Lgr5* expression in mice describes *Lgr5*^+^ mesenchymal cells which are nearly absent from the airways but centered around the alveolar progenitor cells orchestrating the alveolar niche (9). In humans, using scRNA analysis, *LGR5*^+^ mesenchymal cells were identified as airway-associated fibroblasts (PBF) (10,11). The disconnect between human and murine findings reinforces the need for an animal model that represents a more faithful anatomical and physiological model to the human lung (12–17) and can be used to study the function of lung LGR5^+^ cells.

Here we used a validated transgenic porcine model expressing a nuclear-localized histone H2B–GFP under the control of the endogenous *LGR5* promoter (18). This model faithfully identifies LGR5 cells, and we have shown that the H2B-GFP expression correlates spatially and quantitatively with *LGR5*^+^ mRNA expression by RNA in situ hybridization and quantitative PCR (18). Using this model, we show that in pig lungs there are two populations of LGR5^+^ cells in homeostasis and development. We moreover define the spatial organization and transcriptional signature of both populations and compare them to equivalent adult and fetal human cell populations. Results presented here originated from the PhD thesis of Kathryn Polkoff (19).

## Materials and Methods

### Transgenic porcine and human tissue

Experiments were carried out in accordance with the Institutional Animal Care and Use Committee of North Carolina State University. We used lung tissue from 14 postnatal pigs, five fetal GD80 fetuses, five fetal GD50 fetuses. Data are representative of both sexes. Human tissues were acquired from the Marisco Lung Institute at the University of North Carolina at Chapel Hill or from Duke Biorepository & Precision Pathology Center (BRPC). All research using this human fetal lung tissue was approved by the respective IRB Boards.

### Tissue Clearing

We used the BoneClear procedure (20) with adjustments; perfusion, decalcification, and agarose embedding steps were not conducted, and immunostaining occurred over 7 days with anti-GFP primary antibody (Aves Labs, GFP-1010), and Cy3 donkey anti-Chicken IgG (Jackson ImmunoResearch Labs, 703165155).

### Immunofluorescence

Tissue was fixed in 4% paraformaldehyde at 4°C overnight, dehydrated and embedded in OCT. Organoids were fixed in 2-4% paraformaldehyde and embedded in OCT or stained and imaged as whole mounts.

### RNAscope *in situ* hybridization

RNAscope was performed according to the manufacturer’s instructions (ACD Bio). The tissue was hybridized with custom probes targeting Hs-*SCGB3A2* (544951) and Hs-*LGR5* (311021-C2). A positive control probe against human (139-989 region) cyclophilin B or as a negative control, a probe targeting the bacterial gene dapB were used.

### Quantification of SOX2/SOX9

Quantification was performed using ImageJ (NIH). Statistical analysis comparing ratios of co-expressing cells between GD50 and GD80 sections was performed using a Welch Two Sample T-test in R, *n=*10 photos per group, representative of 2 different fetal lungs each.

### Cell Isolation and Sorting

Samples were minced and digested at 37°C in an enzymatic digest buffer containing 5 mg/mL Dispase (Invitrogen), 2 mg/mL DNase I (STEMCELL Technologies), and 5 mg/mL (adult) or 2mg/mL (fetal) Collagenase Type II (Sigma).

### Organoid Culture

2 x 10^3^ cells/droplet were mixed with 25 μl of Matrigel and allowed to solidify upside down at 37℃ for 20 minutes and then overlaid with culture media, replaced every other day.

### RNA sequencing analysis

For bulk RNA-sequencing, sorted mesenchymal cells were used for Ultra-low input RNA-seq. The trimmed reads were mapped to the Sus scrofa reference genome ENSEMBLv11.1.109 assembly. Using DESeq2 (v1.37.6). The Wald test was used to generate p-values and Log2 fold changes. Genes with adjusted p-values < 0.05 and absolute log2 fold changes > 1 were called as differentially expressed genes for each comparison.

### Single cell RNA sequencing

scRNAseq libraries from sorted GD80 LGR5^+^ fetuses were prepared using the 10X genomics system. After sequencing, barcodes were deconvoluted into a cells x genes matrix using alevin v1.5.2 based on the Sus scrofa Ensembl v11.1.105 assembly, and counts were imported into R (v4.2.0) for analysis with the Seurat package (v4.1.1) using the RStudio IDE (v2022.07.1). For quality control, cells expressing less than 2000 counts RNA or less than 1000 genes were excluded from the analysis. The remaining 10,482 cells, with an average of 15,855 gene transcripts per cell, were retained for analysis.

## Results

### Distinct expression of LGR5 by epithelial fetal lung bud tips and airway mesenchymal cells

Lung tissue from the *LGR5-H2B-GFP* line co-stained with EpCAM for epithelial cells and vimentin (VIM) for mesenchymal cells shows that in the fetal lung two LGR5^+^ populations exist: one mesenchymal, concentrated in the peribronchial area surrounding the airways, and one epithelial, in the developing bud tips (Figure 1A-J). Mesenchymal LGR5^+^ cells surround both proximal and developing distal airways but are absent in bud tips (Figure 1D-J). In contrast, airway epithelial cells are LGR5^-^ except at the bud tips (Figure 1H-J). In the postnatal lung, the peribronchial mesenchymal population surrounding the airways remains LGR5^+^, but all epithelial cells are LGR5^-^ (Figure 1K-S). To determine whether the distribution of LGR5^+^ cells in the pig lung is representative of the human, we employed fluorescent *in situ* hybridization (FISH) to detect *LGR5* mRNA transcripts in human fetal and adult lung tissue. *LGR5* expression is robust in human fetal lung bud tips (Figure 1T). *LGR5*^+^ expression was also detected in the subepithelial population surrounding maturing airways (Figure 1U). Finally, robust detection of *LGR5* is evident in the peribronchial region of adult airways (Figure 1V). For three-dimensional spatial mapping of the LGR5^+^ cells, we used tissue clearing (19, 20) and light sheet microscopy on gestational day 80 (GD80) and juvenile porcine lungs and confirmed that the mesenchymal cells surround the airways and that epithelial LGR5^+^ cells are concentrated in the elongating buds tips of the developing fetus (Supplement Video). These data support that there are at least two distinct LGR5^+^ populations in the lung, and that their presence is consistent in humans and pigs.

**Figure 1.**
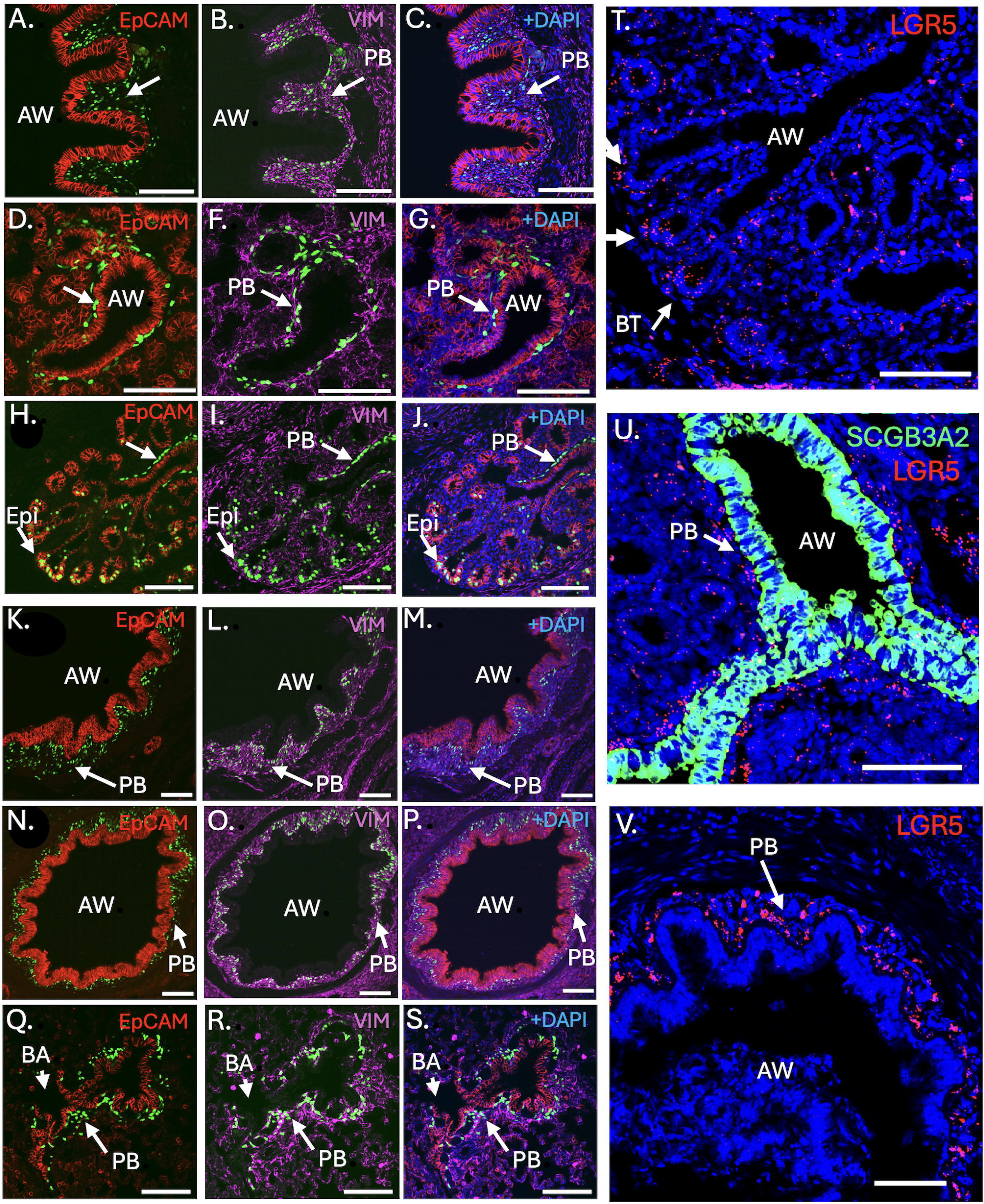
Two distinct populations of LGR5-expressing cells exist in the pig and human lung. **(A-J)** In the GD80 lung, two LGR5^+^ populations are seen, an EpCAM^+^ (red) and a VIM^+^ (purple) one. The population surrounding the airways is mesenchymal (VIM^+^), is located in the developing peribronchiolar region (white arrow, PB), and surrounds developing large proximal (A-C), mid-proximal (D-G) and distal airways (H-J). The LGR5 cells in the bud tips, in contrast, are EpCAM^+^ epithelial cells (white arrow, Epi). **(K-S)** In the postnatal mature pig lung, the VIM^+^ mesenchymal population is present in airways of all sizes (K-M, proximal; N-P, mid-proximal; Q-S, distal) up to the bronchioalveolar region (BA), LGR5^+^ cells are in the sub-basal or peribronchial (PB) space, and no LGR5^+^ epithelial cells are detected. **(T-V)** Conservation of *LGR5*^+^ populations between humans and pigs. *In situ* hybridization with a probe detecting human *LGR5* transcripts in human developing lung shows the presence of epithelial *LGR5* expression in gestational day 135 bud tips (T, white arrows, BT), and the presence of sub-basal/peribronchial LGR*5* expression in cells directly adjacent to the airway epithelium (stained with SCGB3A2) in both fetal day 142 (U) and adult (K) airways. (AW-airway) Scale bars are equivalent to 100 μM.

### Expression patterns of LGR5+ cells in the pig developing lung bud tips

We examined at the pseudoglandular (GD50) and canalicular stage (GD80) the co-expression of LGR5 with markers of bud tips, including SOX9 and SFTPC (21), as well as a marker for more developed airway cells, SOX2 (22). LGR5 cells co-express SOX9 (LGR5^+^/SOX9^+^) and locate to the bud tips intercalated with LGR5^-^/SOX9^+^ cells (Figure 2 and Figure S1). Quantification of the proportion of LGR5^+^ cells that co-express SOX9 shows that at GD50, 42% SOX9 cells are also LGR5. This decreases to 22.2% by GD80 (P<0.05). LGR5 expression is also seen in low-expressing SOX2 cells, and they decreased from GD50 (40%) to undetectable at GD80 (Figure 1S) (P<0.05). Combined, these results suggest that epithelial LGR5 expression is increasingly restricted as the cells transition from SOX9 to SOX2 expression (tip versus stalk) (19). Moreover, *RSPO2*, a known ligand of LGR5, is detectable in the stromal region surrounding the bud tip near LGR5^+^ cells at both GD50 (Figure 2J and 2L) and GD80 (Figure 2K and 2M). LGR5 expression is also correlated with epithelial SFTPC expression (Figure 2N-O).

**Figure 2.**
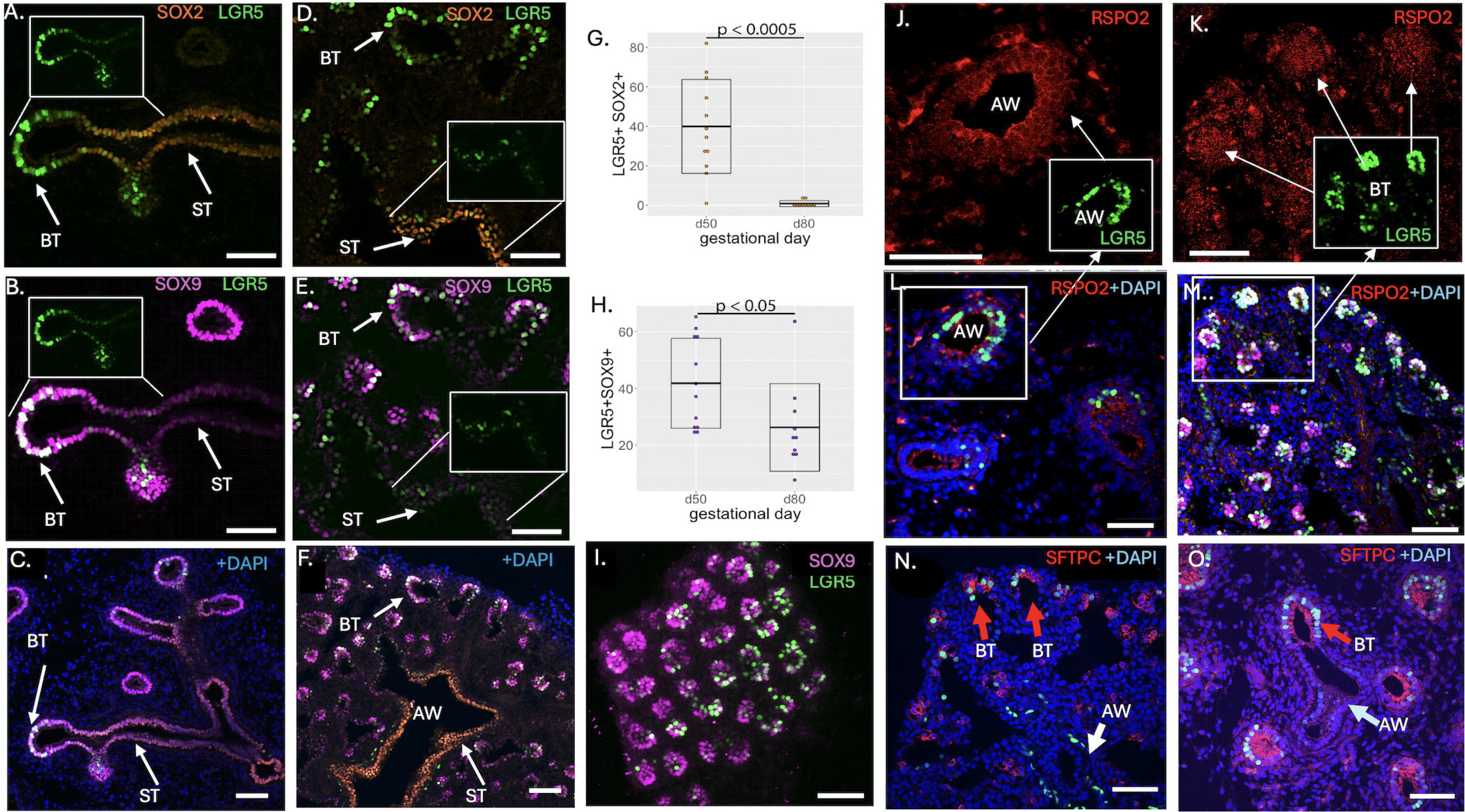
Epithelial LGR5^+^ cells are spatially located in the distal portion of elongating lung bud tips and become more distally restricted as development progresses. (A-F) Representative images of the relationship between LGR5 and SOX9, a known marker of developing bud tips, and SOX2, a marker of more differentiated airways. (G) Quantification of LGR5 and SOX2 co-expression, represented as percentage of co-expressing cells over total LGR5^+^ cells. co-expression is significantly reduced between GD50 and GD80 (p<0.0005, Welch two sample t-test, n=10 images representative of sections from 2 different fetal lungs each). (G-H) Percentage of LGR5^+^ as a proportion of total SOX9^+^ cells decreases from GD50 to GD80 (p<0.05, Welch two sample t-test n=10 images, 2 different fetal lungs each). (I) 3D visualization of LGR5 and SOX9 expression at GD80 via whole-mount confocal imaging showing intercalated SOX9 and LGR5 expression in lung bud tips. (J-M) Expression of RSPO2 in the region surrounding bud tips (white arrows) at GD50 (J, L) and GD80 (K, M). (N-O) Fetal lungs showing co-expression of LGR5 and the known lung bud tip marker SFTPC (BT, red arrow) at GD50 (N) and GD80 (O). (BT, red arrow). (AW-airway) Scale bars are equivalent to 100 μM.

### High-resolution transcriptome analysis of fetal lung LGR5+ cells

Fluorescence-activated cell sorted (FACS) fetal lung LGR5^+^ cells from GD80, a stage where both mesenchymal and epithelial cell populations coexist, were used for scRNA-Seq. We identified a total of four epithelial (*EPCAM*^+^) and ten mesenchymal (*VIM*^+^) clusters and calculated the percentage of cells (%) within each cluster in relation to the whole population (Figures S2-S3). Of the 14 clusters, four are proliferating: Epi 4 (0.3%), Mes 5 (9.3%), Mes 8 (2.4%) and Mes 9 (1.4%); Figure S2C-D). Using Monocle3 pseudotime analysis we show a predicted order of epithelial clusters as Epi 3 (2.0%), Epi 1 (6.3%), Epi 2 (4.1%) with Epi 4 being a proliferating cluster (P) (Figure 3A). This trajectory is supported by proximal-distal markers; Epi 1 and Epi 3 express distal airway progenitor cell markers *SFTPC, SFTPB, ETV5, SOX6*, and *SOX9*. Epi 2 is defined as more proximal based on expression of *KLF5* and *THRB* (Figure 3B). We also examined markers of postnatal distal immature AT1 cells described in humans (11) including *AGER, HHIP, RNASE1*, and *SFTPB* (Figure 3B). In addition, epithelial cell-cell signaling pathways known to be involved in lung development were examined. *BDNF* receptors *F11R* and *SORT1* were expressed by Epi 1 and Epi 3, respectively. All Epi clusters also express *FGFR2* and *FGFR3*, receptors for *FGF9* and *FGF10*, and *FZD2/3/5* receptors for *WNT2* and *WNT2b*. Levels of *SHH* increase in order of cluster Epi 3, Epi 1 and Epi 2 (Figure 3C). A more comprehensive analysis of *SHH, FGF, TGF-*β and *WNT* pathways genes are shown to further support the distal/proximal arrangement of the epithelial clusters (Figure S4).

**Figure 3.**
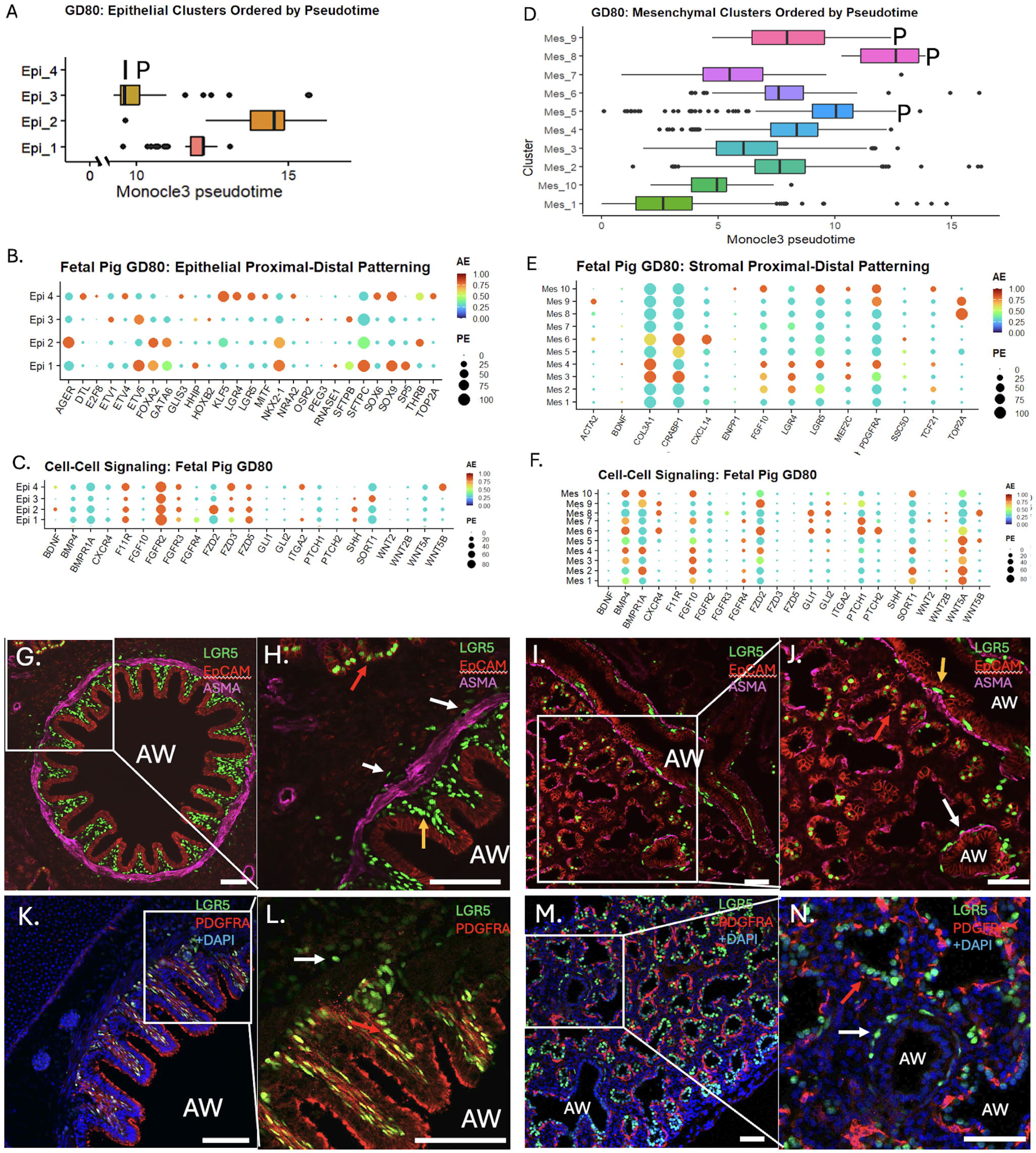
Single cell RNA sequencing of GD80 fetal porcine LGR5^+^ cells identify distal-proximal cell types and active key signaling pathways in both epithelial and mesenchymal clusters. (A) Pseudotime analysis shows a predicted order of epithelial clusters as Epi 3, Epi 1, Epi 2, with Epi 4 being a proliferating cluster (P). (B) Epithelial markers associated with proximal-distal patterning of the elongating lung bud including *ETV5, SFTPC, SOX6* and *SOX9*. (C) Epithelial cell-cell signaling pathways known to be involved in lung development. Both ligands and their receptors are shown. (D) Pseudotime analysis showing a complex mesenchymal cluster trajectory. (E) For the mesenchymal clusters, multiple known markers that define mesenchymal subsets, including those defined by Cao et al (23), were used. They include *BDNF*, *CRABP1*, *COL9A9* (mapped using *TCG21,* an alternate marker for this population), *ACTA*, and *TOP2A*, and *PDGFRA*. (F) Cell signaling by clusters was examined via receptors and ligands known to be involved in lung development and function. (G-H) Staining of GD80 proximal airway with ASMA and EpCAM, shows that most of the LGR5^+^ mesenchymal cells reside within the peribronchial region (H-yellow arrow). However, LGR5 cells are also seen distal to the airways (AW) and the smooth muscle cells (H-white arrows). In addition, at GD80 LGR5^+^ epithelial cells can still be observed (H-red arrow). (I-J) GD80 distal immature airway (I-box and J-white arrow) with mesenchymal LGR5^+^ distal to a small layer of smooth muscle cells. In contrast, in the more developed airway (J-orange arrow) peribronchial LGR5^+^ cells are observed. (K-L) Proximal airway showing peribronchial LGR5^+^/PDFGRA^+^ (L-red arrow) but also the presence of LGR5^+^/PDGFRA^-^ cells (L-white arrow). (M-N). In the distal lung, immature airways are surrounded by PDGFRA^-^/LGR5^+^ mesenchymal cells (M-Box and N-white arrow). Note the presence of PDGFRA^+^/LGR5^-^ (N-Red arrow). Scale bars are equivalent to 100 μM. AE: average expression; PE: percent expression.

For mesenchymal clusters, pseudotime analysis showed a complex trajectory. Mes 1 (20.3%), a *PDGFRA^-^* cluster is mapped at one end of the trajectory while the three proliferating clusters Mes 5 (9.3%), Mes 8 (2.4%) and Mes 9 (1.4%) are at the opposite end. The remaining clusters have significant overlap with no clear order suggesting either two independent lineages and/or a branched lineage. We also examined whether the stromal subsets defined by Cao et al (23), *SC_BDNF*^+^, *SC_CRABP1*^+^, *SC_COL9A*^+^ (mapped using *TCG21*, an alternate maker for this population), SC*_ACTA*^+^, and *SC_TOP2A*^+^, were represented in the different clusters. No obvious *SC_BDNF*^+^ cluster was detected with all clusters expressing low levels of BDNF. Other clusters could be mapped to these subsets, *SC_CRABP1*^+^ [Mes 3 (12.5%) and Mes 6 (8.6%)], *SC_COL9A*^+^ [Mes 2 (14.3%), Mes 4 (10.7%), and Mes 10 (0.7%)], *SC_ACTA2*^+^ (Mes 6), and proliferating populations *SC_TOP2A*^+^ (Mes 5, Mes 8 and Mes 9) (Figure 3E). We also mapped non-proliferating *PDGFRA*^-^ clusters (Mes 1 and Mes 7) (Figure 3E). Mesenchymal cell-cell signaling showed expression of ligands and receptors (Figure 3F). Ligands include low levels of *WNT2* (Mes 7) and *WNT2B* (Mes 2 and Mes 7), and low levels of *BMP4*, *FGF10* and *WNT5A*. Receptors include *BMPR1A, FGFR2*, and *SORT1*. In addition, several genes known to be activated by SHH are expressed by all mesenchymal (Mes) clusters (Figure 3F). A more comprehensive signaling signature of all mesenchymal clusters is presented in Figure S5.

To spatially anchor the different clusters, we examined the expression of ASMA, EpCAM and PDGFRA (Figure 3G-N). GD80 proximal airway shows that most of the EpCAM^-^/LGR5^+^ mesenchymal cells reside within the peribronchial region (Figure 1H). However, EpCAM^-^/LGR5^+^ cells are also seen distal to the airways (AW) and the smooth muscle cells outside of the peribronchial region (Figure 3H). In the GD80 immature airway, EpCAM^-^/LGR5^+^cells can be seen distal to a small layer of smooth muscle cells (Figure 3J). In the more developed distal airway (Figure 3J), a few peribronchial EpCAM^-^/LGR5^+^ cells are observed. These data support the existence of non-peribronchial LGR5^+^ mesenchymal cells in both the mature proximal and the developing distal airways. We also examined PDGFRA expression to determine if we could spatially map the non-replicating PDGFRA^-^ clusters (Mes 1 and Mes 7) and detected two PDGFRA^-^/LGR5^+^ populations, one located outside the smooth muscle layer in the more developed airways (Figure 3H) and one in the distal immature airways (Figure 3N). Attempts to map the different LGR5^+^ GD80 clusters to specific cell populations using GSEA failed, likely due to insufficient fetal lung data within GSEA.

### Functional analysis of fetal and adult epithelial cells via organoid modeling

As the expression of LGR5 is increasingly restricted in the epithelial population between GD50 and GD80, we asked whether LGR5 expression and developmental stage impacted their ability to form organoids (19). LGR5^+^ or LGR5^-^ epithelial cells were sorted from fetal GD50 or GD80 lungs and plated in 3D culture. LGR5^+^ cells from GD50 lungs formed significantly more organoids than any other group, while there was low organoid formation from GD80 (Figure 4A). Overall, this suggests changes in cell state during development, where the LGR5^+^ bud tip cells respond differently to inductive signals depending on stage of development. In addition, a few organoids from postnatal airway epithelial cells (LGR5^-^) seeded under the same conditions displayed detectable LGR5^+^ cells at 5-7 days, and by day ten, most, if not all, organoids had LGR5^+^ cells (Figure 4B). We next asked if there were any differences between the behavior of organoid-derived LGR5^+^ and LGR5^-^ cells. After dissociation, both populations were sorted (Figure 4C) and re-seeded separately. We observed no detectable difference between the two populations in organoid forming efficiency, and organoids from the LGR5^-^ population once again reactivated LGR5 (Figure 4C).

**Figure 4.**
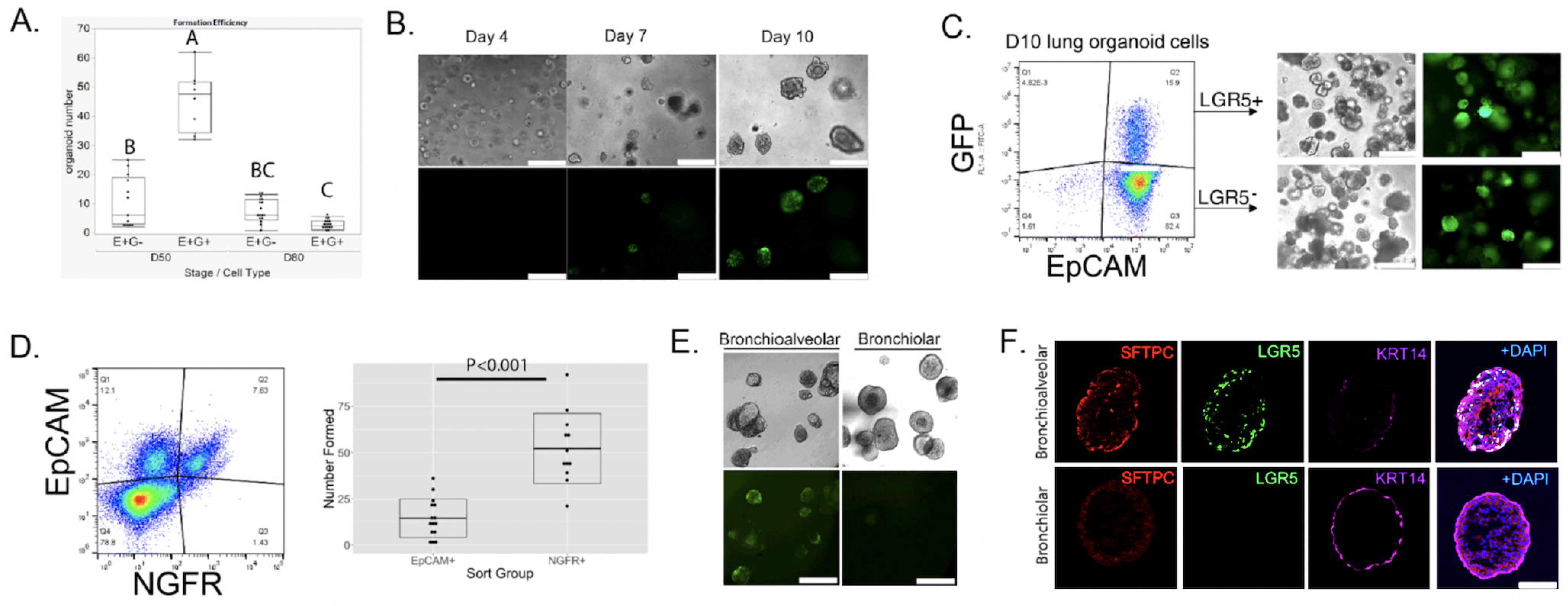
LGR5 expression is transiently reactivated in adult epithelial organoids and is associated with organoid forming efficiency and cell fate. (A) Organoid formation efficiency of EpCAM^+^/LGR5^+^ (E^+^G^+^) and EpCAM^+^/LGR5^-^ (E^+^G-) cells from GD50 or GD80 fetal pigs with GD50 EpCAM^+^/LGR5^+^ cells demonstrating the highest formation efficiency. Comparison of all pairs was performed using a one-way ANOVA by group, and comparison of means was performed by Tukey-Kramer HSD, n=3 biological and 10 technical replicates for each group, p<0.05 (B) EpCAM^+^/LGR5^-^ cells from adult lungs were placed in organoid culture. While organoids remained LGR5^-^ after 4 days of culture, by 5-7 days LGR5 expression was detectable. By day 10, a mixture of LGR5^+^ and LGR5^-^ organoid could be detected. Scale bars represent 300 uM. (C) After 10 days in culture, organoids were dissociated and sorted by EpCAM and LGR5 (GFP) expression. EpCAM^+^/LGR5^-^ and EpCAM^+^/LGR5^+^ cells were plated separately for organoid formation, and both populations generated a mixture of LGR5 positive and negative organoids. Scale bar represents 300 uM. (D) LGR5^+^ adult lung cells were sorted based on expression of EpCAM and NGFR (an airway basal cell marker) and EpCAM^+^/NGFR^+^ and EpCAM^+^/NGFR^-^ cells were plated separately for organoid formation efficiency. EpCAM^+^/NGFR^+^ epithelial cells formed organoids at a significantly higher efficiency than EpCAM^+^/NGFR^-^ cells (p<0.001, Welch two sample t-test, n=3 biological and 4 technical replicates each). (E) Postnatal epithelial cells (LGR5^-^) were cultured in bronchioalveolar (BA) or bronchiolar (BR) organoid media and imaged by fluorescent or brightfield microscopy. Only bronchioalveolar culture conditions induced LGR5 expression. Scale bar represents 300 uM. (F) Organoid type: bronchiolar (BA) or bronchioalveolar (BR) was confirmed by morphology and by IHC using antibodies to SFTPC and KRT14. Scale bars represent 900 uM. AE: average expression; PE: percent expression.

To ask which cells reactivated LGR5 expression and formed organoids, we plated sorted lung cells based on NGFR, a known basal cell marker, and EpCAM (Figure 4D). The EpCAM^+^/NGFR^+^ basal cells formed significantly more organoids than the EpCAM^+^/NGFR^-^ cells, but both expressed LGR5 and had mixed LGR5^+^ and LGR5^-^ organoids, suggesting that, although NGFR^+^ basal cells were the primary driver of organoid formation, the basal cells and non-basal cells had similar fate under these conditions (19).

This observation led us to ask what the cell state of the organoid LGR5^+^ versus LGR5^-^ cells is by RNA-seq analysis (Figure S6A). Among the significantly upregulated genes in the LGR5^+^ population were *SFTPC, KRT14, AXIN2*, markers of specialized progenitor basal and AT2 cells (24). In addition, LGR5^+^ cells upregulated *BMP4*, *BMPR1B* and *TGFB2*, all known regulators of lung airway development (25) and expressed by GD80 epithelial cells (Figure S4). Among the genes downregulated in LGR5^+^ cells were *AQP5*, an alveolar AT1 cell marker (26) *LTF*, a marker for differentiating basal cells (27) and *FRZB, FZD6* and *FST*, known negative regulators of canonical WNT signaling (26,28–30).

Given the shared markers of LGR5^+^ organoid cells with lung development markers, we hypothesized that the LGR5 expression was related to cell fate. To test this, we altered the media conditions from a bronchioalveolar (BA) fate to a bronchiolar (BR) phenotype by reducing WNT and WNT agonist CHIR (19).

After ten days in culture, none of the bronchiolar organoids expressed LGR5, while the bronchioalveolar were LGR5^+^ (Figure 4E). To classify the phenotype of each condition, we confirmed the expression of KRT14 (basal cell marker) and SFTPC (31) (Figure 4F). These data suggest LGR5 is only expressed when airway epithelial cells are differentiating toward an alveolar fate, but not when they maintain a basal cell state.

### Adult LGR5+ mesenchymal cells form a unique niche in the peribronchial region and express a stromal cell gene signature

As myofibroblasts are a known mesenchymal population surrounding airways, we asked if LGR5^+^ cells in the adult lung were ASMA^+^ (αSMA^+^) myofibroblasts. As shown in Figure 5A-F, LGR5^+^ and ASMA^+^ populations remain separate; in proximal airways the LGR5^+^ cells are concentrated between the ASMA cells and the airway epithelial cells (Figure 5B-C), and in the distal bronchioalveolar region the two populations are intercalated (Figure 5E-F), supporting separate roles for ASMA and LGR5 populations. Other minor LGR5^+^ populations, also ASMA^-^, can be seen distal to the smooth muscle layer (Figure 5D-F, purple arrow) or in the alveolar region (Figure 5D, green arrow) (19). The distinct nature of the ASMA^+^ and LGR5^+^ populations is supported by a 3D image of a proximal airway showing they are arranged in perpendicular orientation to each other (Figure 5G-H). In addition, peribronchial (PB) LGR5^+^ cells are located directly adjacent to the NGFR^+^ basal cells (BC) (Figure 5I). Finally, at GD80, a stage that encompasses well developed to immature airways, the distribution of the LGR5^+^ cells differed with the degree of spatial organization both longitudinally and circumferentially decreasing proximally to distally (Figure 5J-L).

**Figure 5.**
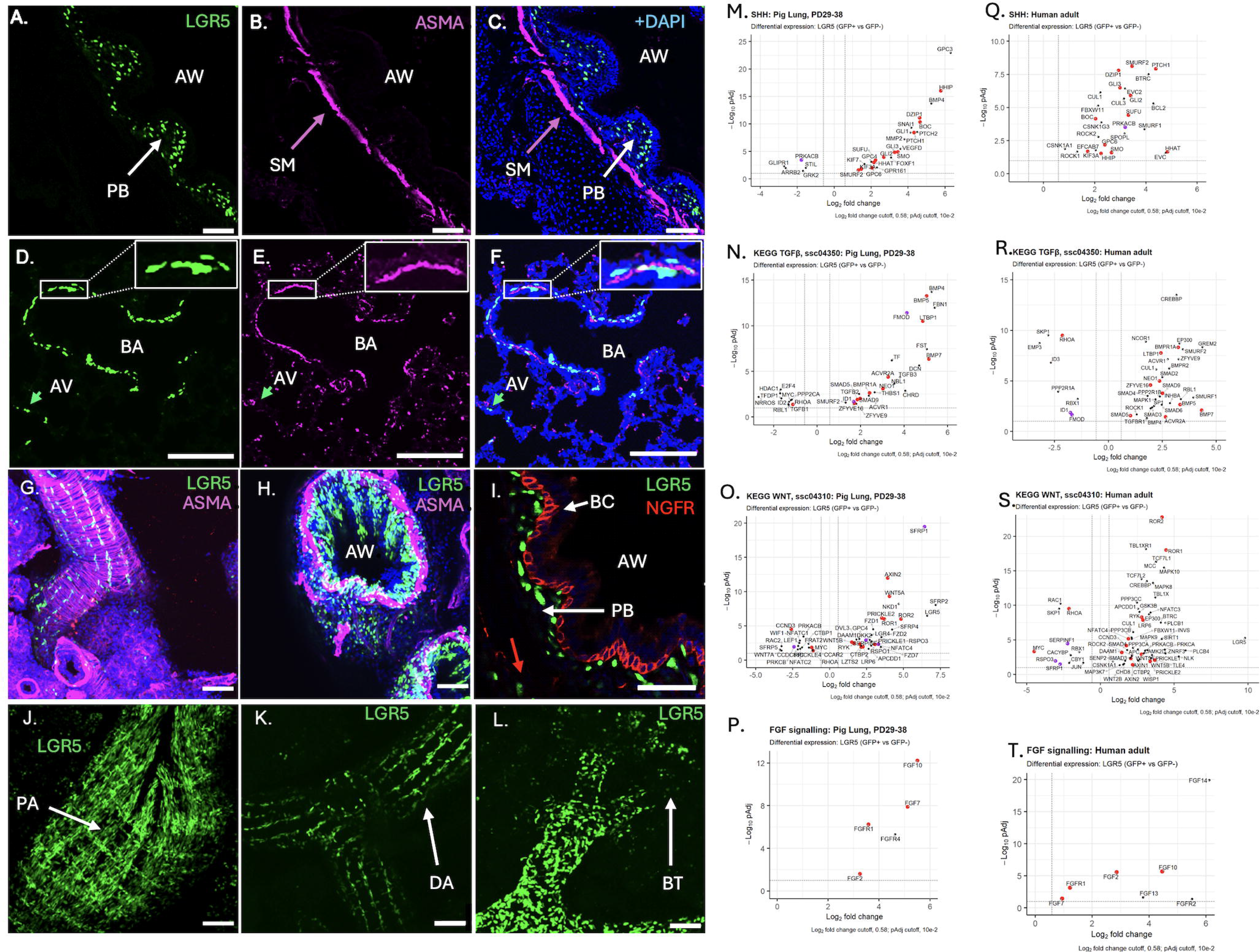
LGR5^+^ mesenchymal populations are distinct from myofibroblasts, have a highly organized spatial distribution in the mature airways, lie in proximity to airway basal cells and have a unique transcriptional profile. (A-F) Confocal imaging of bronchiolar and bronchioalveolar cross-sections, shows that, like GD80 fetuses, mesenchymal LGR5^+^ cells are spatially distinct from ASMA expressing smooth muscle and myofibroblasts. (A-C) In the mature airways LGR5^+^ cells are concentrated in the peribronchial space (PB-white arrow) between the airway epithelia and the ASMA^+^ layer (SM-Purple arrow) and the lung airway epithelial cells. (D-F) In the bronchioalveolar region (BA), LGR5^+^ and ASMA^+^ cells are separate, and can be seen in both sides of the ASMA layer (SM-Purple arrow). In addition, some ASMA^-^/LGR5^+^ cells can be seen in the alveolar region (AV-green arrow) (G-I) 3D rendering of whole mount tissue shows the longitudinal arrangement of the LGR5^+^ cells perpendicular to the orientation of the ASMA^+^ cells. (I) Close apposition between LGR5^+^ cells and the basal airway epithelial cells (BC) identified by NGFR expression. Note also non-peribronchial LGR5^+^ cells (red arrow). (J-L) Spatial organization in GD80 whole mount confocal images, with both longitudinal and circumferential alignment in large proximal airways (PA), primarily longitudinal alignment in smaller airways (DA), and more random organization in distal airways near the elongating bud tips (BT). Scale bar indicates 100 μM unless marked otherwise. (M-P) SHH, TGF-β, WNT, and FGF pathways genes that are differentially expressed in postnatal (PD29-PD38) LGR5^+^ versus LGR5^-^ mesenchymal cells. (Q-T) SHH, TGF-β, WNT, and FGF pathway genes are differentially expressed in human LGR5^+^ versus LGR5^-^ mesenchymal cells. For these volcano plots, LGR5^+^ and LGR5- cells were extracted from the peribronchial fibroblast cluster in the human lung dataset published by Madissoon et al (10). Genes differentially expressed in both pigs and humans (concordant) are shown in red.

As the LGR5^+^ cells occupy a niche directly adjacent to the airway basal cells, and the GD80 data indicated fetal LGR5 cells had a complex *SHH, FGF,* and *TGF*β signaling, we asked whether these pathways were significantly enriched in the adult LGR5^+^ population by comparing the transcriptome of LGR5^+^ and LGR5^-^ postnatal mesenchymal cells (Figure 5M-P). Adult LGR5^+^ cells upregulated key *SHH* pathway genes such as *SMO, SUFU, PTCH2, GLI1*, and *GLI2*. We also examined *TGF-*β pathway genes as they are known to be involved in lung development and disease (32–34) and interact with the WNT pathway (35,36). As shown in Figure 5N, *TGFB2, TGFB3, BMP4, BMP5* and *BMP7* were upregulated in LGR5^+^ cells. For the WNT pathway, adult LGR5^+^ cells express canonical and non-canonical WNT signals, such as *WNT5A, RYK, ROR1, ROR2, and GPC4*, in addition to known negative regulators of WNT signaling, such as *SFRP1, SFRP2 and PRICKLE2*. Furthermore, they upregulate WNT self-responsive signals such as *FZD1, FZD2* and *AXIN2* (Figure 5M) (37). Finally, we examined FGF signaling due to their known role in lung development and function and detected upregulation of ligands *FGF2, FGF7*, and *FGF10* and the receptors *FGFR1* and *FGFR4* (Figure 5P). Many, if not all, of the genes above were also expressed in one or more GD80 mesenchymal clusters (Figure 3C and Figure S5)

Next, we compared pig and human LGR5^+^ peribronchial fibroblasts transcriptome by analysis of the Madissoon et (10) scRNAseq data. We extracted all 729 *LGR5*^+^ cells and mapped them to the human lung cell atlas. As shown in Figure S7, *LGR5*^+^ cells were concentrated in the peribronchial fibroblasts cluster (Oval), with smaller populations in other clusters. Using pathway scores generated from data shown in Figure 5M-T we also mapped the *SHH, TGF*β*, FGF*, and the canonical and non-canonical (polar cell polarity/PCP) pathways. While the cell signatures scores mapped to more than one population, comparison of the different stromal subsets shows that the dominant population in all cases is the peribronchial population. Overall, this supports that the signaling properties of the *LGR5*^+^ cells are unique and are conserved between humans and pigs.

Previously, Murthy et al (11) described a distal airway *LGR5*^+^ mesenchymal cell, with a transcriptional profile, that was distinct from adventitial fibroblasts, alveolar fibroblasts, fibromyocytes, airway smooth muscle cells, vascular smooth muscle cells, and pericytes. Mapping of this profile to the pig GD80 clusters showed that *WNT5A*, *PAPPA, ASPN, and DPT* were expressed by one or more Mes clusters. Equally important genes not expressed in *LGR5*^+^ fibroblasts, including *WNT2, DES, MYH11, CNN1* and *RAMP1* were also not expressed at high levels in any of the clusters, and genes expressed by *LGR5^+^,* as well as adventitial, and alveolar fibroblasts including *CCDC80, SELENOP, FN1, C7, RGCC, MACF1, LUM* and *GPC3* were expressed by all clusters.

To further characterize the pig LGR5^+^ cells we applied Gene Set Enrichment Analysis (GSEA) (38) to the top 500 upregulated genes in LGR5^+^ PD29-38 mesenchymal cells (Data Table S1) and examined cell type (Module C8), Gene Ontology:Biological Process, and Gene Ontology:Molecular function. As shown in Table S1A, the LGR5^+^ cells have a gene signature that matches adult and fetal stromal cells from multiple organs (19). Additionally, despite being an adult cell, the LGR5^+^ transcriptional signature mapped to fetal cells, and GO-Biological Processes related to development (Table S1B). In terms of molecular function, LGR5^+^ cells are enriched for extracellular matrix constituents and binding activity (Table S1C).

### Adult lung mesenchymal LGR5+ cells serve as a signaling niche for airway basal cells

Because stromal cells can define a support niche for stem cell subsets (11), we hypothesized that the LGR5^+^ cells designate the support niche for airway epithelium. To test this, we isolated NGFR^+^ basal cells from lung airways and co-cultured at a 1:1 ratio with LGR5^+^ mesenchymal cells with or without growth factors and show that LGR5^+^ cells alone could support organoid formation from basal cells Figure 6). Furthermore, organoids in the LGR5^+^ co-culture wells, and growth factor only wells, differed morphologically. Quantification of organoid type as previously described (31) shows that the presence of the growth factors induced a bronchioalveolar fate, but LGR5^+^ mesenchymal cells in basic media supported the growth of bronchiolar organoids, and more specifically KRT14^+^ basal cells (Figure 6) (19).

**Figure 6.**
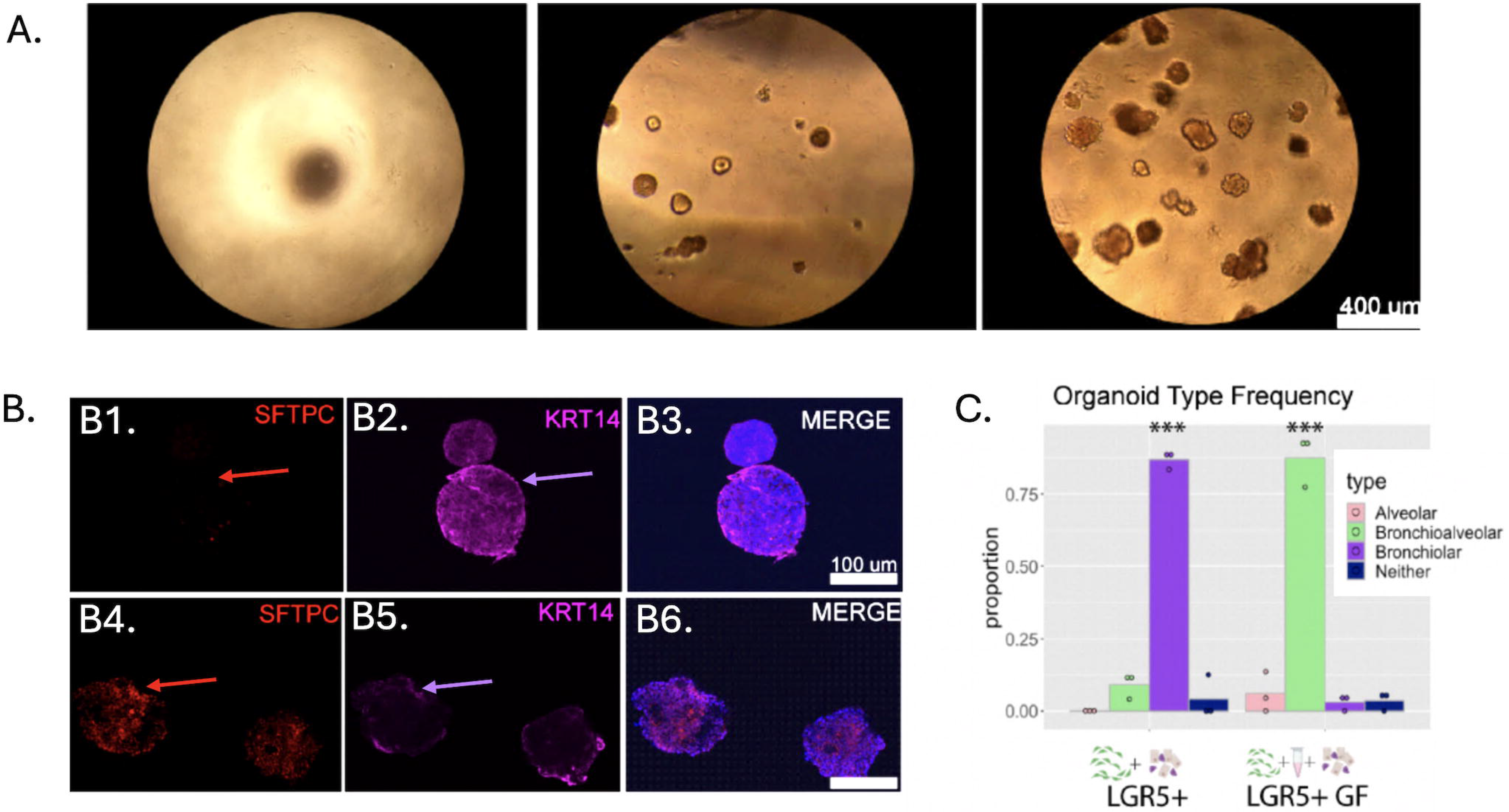
Postnatal LGR5^+^ cells support the airway basal cell niche. (A) Airway basal cells were seeded in growth factor deficient Matrigel either with no additional factors, in a 1:1 ratio with LGR5^+^ mesenchymal cells only, or with a 1:1 ratio of LGR5^+^ mesenchymal cells in a growth factor cocktail. (B1-B3) Bronchiolar organoids were characterized by a spherical shape, absence of SFTPC (B1-red arrow) and robust expression of KRT14 (B2-purple arrow). (B4-B6) Bronchioalveolar organoids had an irregular shape and expressed both SFTPC (B4-red arrow) and KRT14 (B5-purple arrow). Representative images are shown. D) Proportion of organoid type for each growth condition. Organoids from epithelial cells co-cultured with LGR5^+^ cells only were primarily bronchiolar, while conditions containing LGR5^+^ cells and growth factors or growth factors alone yielded primarily bronchioalveolar organoids. *** indicates p<0.005, ANOVA for each treatment followed by pairwise t-tests to determine significant differences, points represent n=3 biological replicates which each included at least 25 organoids per group.

### LGR5^+^ peribronchial mesenchymal cells are associated with nerve fibers

Based on the presence of nerve fibers in proximity to the airways, and the linear-like spatial organization of the cells around the airways, we examined the relationship between LGR5^+^ cells and expression of PGP9.5, a pan-neuronal marker that has been shown to label nerve fibers in adult and fetal lungs (39–42), TUJ1, a neuron specific class III β-tubulin, S100B, a pan-astrocyte marker expressed in glial cell precursors (43,44) and NGFR/P75 and TRPV1, expressed in sensory C-fibers (45–48). We show that peribronchial LGR5^+^ mesenchymal cells surrounding the airways are organized around nerve fibers that co-express PGP9.5, S100B, NGFR, TRPV1, and TUJI (Figure 7A-F). We also show that LGR5^+^ cells traverse the smooth muscle layers through PI16 negative gaps (Figure 7G), and that blood vessels adjacent to airways also stain for nerve fibers but are devoid of LGR5^+^ cells (Figure 7H). Analysis of RNA-seq data showed that *TUJ1, TRPV1, S100B* and *NGFR* are not detectable in LGR5^+^ indicating that the staining observed is coming from adjacent cells, not the LGR5^+^ cells.

**Figure 7.**
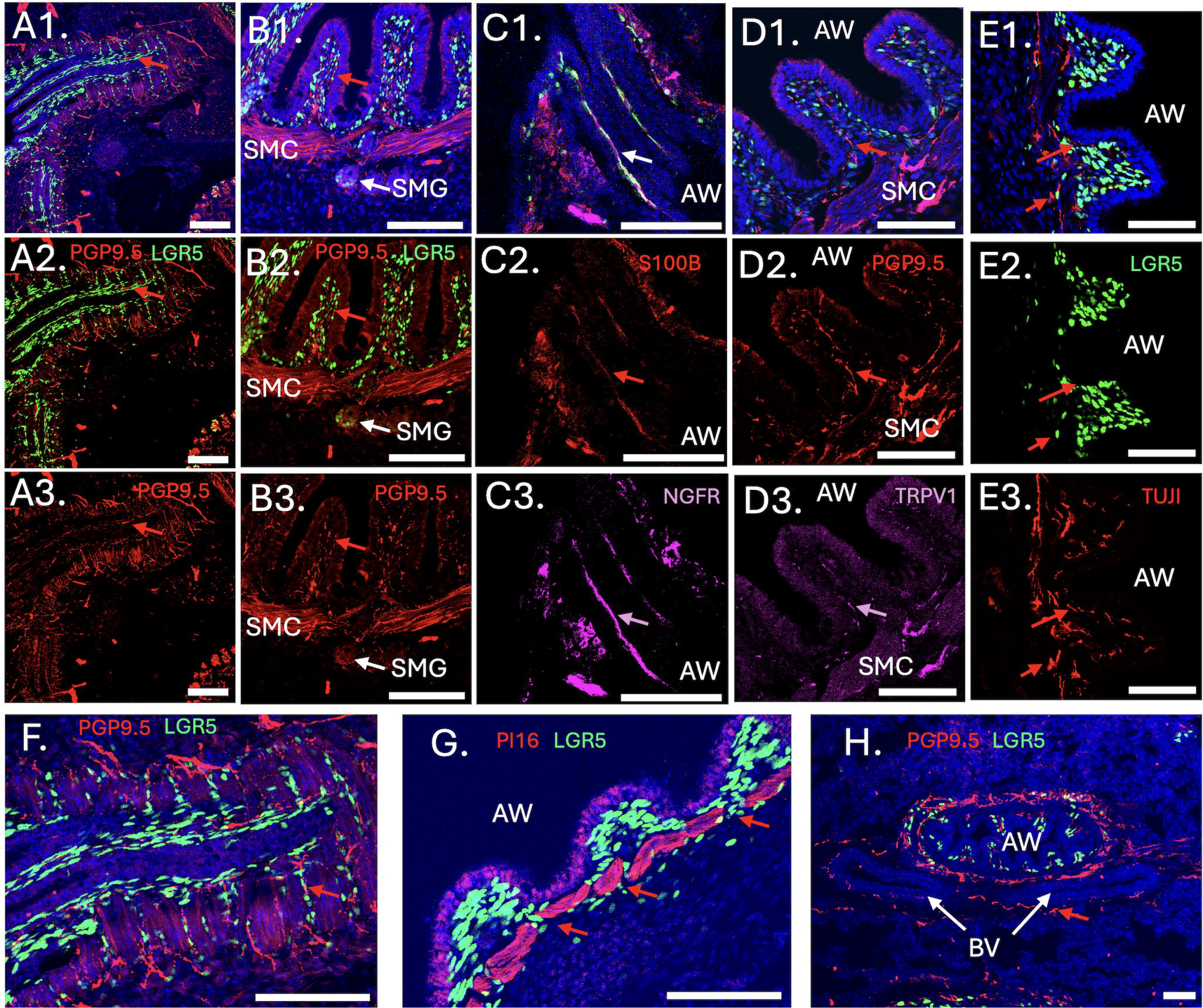
LGR5^+^ mesenchymal cells orient themselves along peripheral nerve fibers. (A1-A3) Sagittal section of airway (AW) with LGR5^+^ cells, identified by their intense nuclear GFP, oriented along PGP9.5 positive fibers. LGR5^+^ nuclei area aligned in an organized pattern with the longitudinal neuronal fibers of the airways (red arrow) (B1-B3) Cross-section of airway showing the spatial adjacency of the PGP9.5 fibers (red arrow) in the peribronchial region as well as distal to the smooth muscle layer. Note smooth muscle cell (MSC) innervation and the presence of weakly staining LGR5 cells in the submucosal glands (SMG). (C1- C3) Proximal airway stained with S100B (red arrow) and NGFR (purple arrow). (D1-D3) Proximal airway stained with PGP9.5 (red arrow) and TRPV1 (purple arrow) showing overlap of the PGP9.5 and TRPV1 signals in the fibers near LGR5+ cells (arrows). (E1-E3) GD80 airway stained with TUJ1 shows nerve fibers adjacent LGR5+ cells (red arrow). (F) Larger magnification of airway stained with PGP9.5 highlighting spatial relationship of LGR5 cells with lateral fibers (red arrow) (G) Distal airway stained with PI16 showing gaps in the smooth muscle layers through which LGR5 translocate to the peribronchial region. (H) Cross-section of postnatal airways show the presence of LGR5^+^ cells in the innervation surrounding the airways but not the blood vessels (BV). Scale bars are equivalent to 100 μM.

Due to its spatial adjacency with the nerve fibers, we hypothesized that the LGR5^+^ cells would share gene signatures with nerve associated fibroblasts (NAF) or Schwann cells (49) and examined lung, sciatic and peripheral nerve scRNA-seq datasets, as well as previously reported canonical markers of nerve associated fibroblasts (10,11,49–55). Using these data, we compiled a set of markers defining the different NAF and Schwann cell types (Table S2) and show that LGR5^+^ cells are enriched for markers associated with NAF but not Schwann cells (Figure S6). We also developed a NAF score and mapped it to the pig GD80, and fetal (56) and adult (10) human lung maps (Figure S8). In pigs, we show that Mes 6, Mes 3 and Mes 4 are enriched for cells expressing NAF markers and in both the fetal and adult maps that *LGR5*^+^ cells overlap with the NAF score. These gene signatures, as well as the spatial adjacency, support that a subset of *LGR5*^+^ mesenchymal cells are specialized NAFs of yet unknown function.

## Discussion

Here, we show that the pig is an excellent model for studying the role LGR5^+^ cells in the lung. The described cell populations, an epithelial and a mesenchymal, are consistent in humans (8,10,11) and pigs, but not consistent in mice (9,57,58). Lgr5 is reported in alveolar fibroblasts in mice, and the roles of the murine Lgr6 cells seem to be more akin to the LGR5 human and pig populations (9), given that they surround and support the airway niche. However, in the mouse the Lgr6 and αSMA populations overlap, whereas they do not in the pig. In addition, scRNA-seq of the mouse lung reported the presence of Lgr5 cells in a population identified as Airway Fibroblast 2 (**59**). However, spatial mapping using transgenic lines indicated that this population is in the alveolar, not the peribronchial region (9). This suggests that, while transcriptionally the mouse Lgr5 cell may be akin to the pig and human peribronchial fibroblast, it is spatially distinct.

We also show that LGR5^+^ cells have a signaling signature marked by active SHH, TGFβ, WNT and FGF pathways (19), and that this signaling signature is conserved in humans. This is supported by Murthy et al (11) that described analogous human mesenchymal population. In fetal development, epithelial LGR5 expression is heterogeneous in bud tip cells in areas of high RSPO2 expression. Their restricted location at the most distal region of the developing lung bud suggests that the epithelial LGR5^+^ cells are an early epithelial progenitor. Human epithelial progenitors have been mapped to the distal tip of the fetal lung by multiple groups. Sountoulidis el at (60) described distal fetal epithelial populations characterized by expression of *SOX9* and a gradual decrease in *ETV5* expression leading to proximal progenitor cells. He et al., (56), described three tip population referred to as Fetal AT1, Fetal AT2, and Early tip characterized by expression of *SOX9, TPPP3, ETV5* and *SFTPC*, and a gradual shift of the tip epithelial progenitors from the pseudoglandular to the canalicular stages accompanied by loss of *TPPP3* expression. At the canalicular stage the tip progenitors are *SOX9^+^/ETV5^+^/SFTPC^+^* and give rise to both Fetal AT1 and Fetal AT2 cells. A similar progenitor trajectory, defined by expression of *SOX9, ETV, SOX2* and novel distal and proximal markers *SOX6, ETV1* and *THRB* were described by Cao et al., (23). In mice, Zepp et al. (61) described analogous early epithelial progenitors referred to as AT2/AT1 cells that give rise to both AT2 progenitors and AT1 progenitors. The AT2/AT1 cells are characterized by expression of *Sftpc/Spock2* and *Hopx*. While different in markers, in both species an early progenitor cell capable of giving rise to both AT2 and AT1 cells has been described. Our results support the existence of a similar population in pigs that, like humans, is also marked by *SOX9^high^/ETV5^high^/SFTPC^high^*(Figure 3B; Epi1 cluster) but is also marked by LGR5. In terms of spatial organization, the intercalated pattern of SOX9^High^, SOX9^medium^, LGR5^High^, LGR5^medium^, and SOX2^Low^ cells at GD50 (Figure S1) matches the canalicular stage model proposed by He et al (55), with tip progenitors (*LGR5^+^/SOX9^+^*) intercalated with fetal AT1, fetal AT2 and stalk cells. In addition, the fetal LGR5 epithelial cells shared some markers with bipotential distal AT1/AT0 epithelial cells described in humans by Murphy et al (11).

Functionally, LGR5^+^ epithelial cells, with their secretion of SHH, expression of FGF receptors and potentiation of the WNT pathway by neighboring RSPO2 (Fig 2J-K), combined with their position in the fetal airway bud tips, suggests that the LGR5 receptor may be important for informing and responding to stromal cells and directing tube elongation, controlling lung airway growth, and maintaining an undifferentiated state as the bud tips elongate throughout development. In the adult, epithelial LGR5^+^ expression is absent, yet it’s reactivated during bronchioalveolar organoid formation. As shown in Figure S6, SFTPC was upregulated in adult organoid LGR5^+^ cells and this was confirmed by IHC (Figure 4F). While there was no differential expression, both ETV5 and SOX9 were expressed by LGR5^+^ and LGR5^-^ epithelial organoids. This supports that organoid formation shifts the adult airway epithelial cells toward a progenitor-like state.

For the mesenchymal LGR5^+^ cells, we identify multiple populations in the adult and developing lung with the predominant being a peribronchial fibroblast population, separate from myofibroblasts (19). Due to its close contact to the basal cells and nerve fibers, and its complex signaling signature, it is positioned to provide a support niche cell for basal cells, and to signal to nerve associated fibroblast involved in communication between the airway cells and the peripheral nervous system. Our RNA-seq data from LGR5^+^ mesenchymal cells show that they have active SHH, TGFβ, FGF and WNT cell-cell signaling. They also express canonical WNT inhibitors and upregulate WNT receptors and downstream WNT pathways. Their unique spatial relationship with basal cells suggests that the LGR5^+^ mesenchymal cell may play an active role in blocking ambient WNT signaling from lung interstitium from reaching the airway. Recent work shows that canonical WNT inhibitors promote basal cell stemness (62), while increased canonical WNT signaling promotes basal cell proliferation in airway repair (63). This suggests that LGR5^+^ cells may influence airway basal cell homeostasis and proliferation by titration of WNT signaling. This interpretation is consistent with that on Murphy et al., (11) that reported human *LGR5^+^* fibroblasts from a signaling hub in the airways niche. And, as we show in Table S3, the *LGR5*^+^ signaling signature described by Murthy et al (11) is consistent with the GD80 mesenchymal clusters.

Our data also supports the existence of non-peribronchial LGR5^+^ cells. As shown in Figure S7, smaller subsets of adult *LGR5*^+^ mesenchymal cells map to the vascular system (blood vessel and endothelial cells) as well as myoepithelial cells and non-peribronchial fibroblasts. Using human fetal atlases (23,56,60) we were able to show that the fetal *LGR5*^+^ cells also have a wide distribution in the human lung with the caveat that the number of *LGR5*^+^ cells in these maps is small. More work is needed to determine the relevance of the close association of LGR5^+^ cells with the nerve fibers. There are reports of innervation impacting airway branching during development (64), epithelial organ regeneration (65) involvement in diseases such as asthma and cystic fibrosis (46,66), and activation of a transcriptional injury signature in airway sensory fibers during inflammation (67). Yet there is little known what role, if any, LGR5^+^ cells play in these responses. And while the LGR5^+^ mesenchymal cells have a signature akin to nerve associated fibroblasts, they appear to be a unique population that is spatially associated with lung airway innervation, but not with the innervation of the adjacent blood vessels, suggesting that they are a specialized nerve-associated fibroblasts, of yet unknown function. Further work is required to understand this complex mesenchymal population and whether it has progenitor properties like the LGR5+ epithelial cells. A weakness of this model is the lack of comprehensive lineage tracing and we are now adding that capability to this model. However, due to the absence of the equivalent cells in mice, this pig model is the only experimental model available at the moment that can be used to examine the role of LGR5 cells in the lung. Overall, this new animal model, and future improved versions, will greatly assist towards the understanding of the role of LGR5 cells during lung development, as well as in the healthy, diseased, and injured lung.

## Data availability

scRNA-seq data and Seurat objects generated in this study are available from the Gene Expression Omnibus (GEO) under accession code GSE218658. The code to process, analyze and visualize the sequencing data is available at https://github.com/RossLampe/Two_roles_of_LGR5_in_the_lung.

The human scRNA-seq datasets used for comparative analysis (54, 65, 66, 67) are publicly available at https://5locationslung.cellgeni.sanger.ac.uk/, https://fetal-lung.cellgeni.sanger.ac.uk/, GSE215895, and OMIX003147, respectively. These data were analyzed using the Seurat package (v5.0.3).

## Author Contributions

KMP, NKG, RL, JAP Conceived and designed the analysis. YM, RL, JC, AC, LE, KG Collected the data.

AM, AG, JMS, PKM, AT, PRT Contributed data or analysis tools.

KMP, RL, YM, JMS Performed the analysis. KMP, NKG, RL, JAP Wrote the paper.

## Acknowledgments

This work was supported by awards F31AR077423 to KMP, T34GM131947 to AC, and R21OD019738 to JAP. Authors are grateful to staff at the Laboratory Animal Resources and the Central Procedures laboratories at the College of Veterinary Medicine, North Carolina State University. The single-cell RNA-seq project was facilitated by the Advanced Analytics Core (partially supported by garnet P30 DK034987) and by the Bioinformatics and Analytics Research Collaborative (BARC) at the University of North Carolina at Chapel Hill School of Medicine.

## Supplement Information

**Supplement Figure S1. Representative images that were used to calculate the co-expression of SOX9, SOX2 and LGR5 to generate the data in Figure 2**. (A-D) GD50 fetus. Note co-expression of SOX2 and SOX9 in most of the developing lung buds. The highlighted areas show co-expression of LGR5, SOX9 and SOX2. Also note that as SOX2 signal intensity increases, SOX9 and LGR5 intensity weakens. (E-H) GD80 fetus. In contrast to GD50, SOX2 co-expression with SOX9 is essentially undetectable even in the tip of the lung buds that are still expressing SOX9 and LGR5. The only SOX2 expressing area (highlighter area) is in the more mature airways now showing surrounding mesenchymal LGR5^+^ cells. The GD80 is at a lower magnification to show a broader area to emphasize the lack of SOX2/SOX9 co-expression. Scale bars are equivalent to 100 μM.

**Supplement Figure S2. Single cell RNA sequencing of GF80 fetal porcine LGR5^+^ cells confirm the presence of epithelial and mesenchymal LGR5^+^ populations during development.** LGR5^+^ cells were FACS sorted from porcine fetal GD80 lungs, and after quality assessment and filtering, 10,482 cells with 15,855 transcriptions per cell were retained for analysis. A) UMAP analysis identified two main populations marked by differential expression of EPCAM and VIM. (B) Examination of LGR5 expression in all clusters showed that mesenchymal populations had higher levels of LGR5 expression than the epithelial populations. (C) Cell cycle markers show mesenchymal clusters 8 and 9 are in G2M. (D) Cell cycle markers show mesenchymal cluster 5 and epithelial cluster 4 are in S phase, and mesenchymal cluster 8 has cells in both G2M and S Phase. AE: average expression; PE: percent expression.

**Supplement Figure 3. GD80 fetal stage reduced set Presto markers and distribution of LGR5+ cells per cluster.** (A) Reduced set of Presto markers used to define the ten mesenchymal (VIM+) sub-clusters and four epithelial sub-clusters (EPCAM+). A full list of markers is found in Supplement Table 1. (B) The percentage of cells in each of the clusters as a total of the whole population (10,482 cells). AE: average expression; PE: percent expression.

**Supplement Figure S4. SHH, FGF, TGF-β and WNT pathway genes are expressed by the different epithelial clusters at GD80. (A)** SHH pathway genes. Note expression of *SHH* by clusters 1-3. **(B)** FGF pathway genes. Note expression of both *FGFR1* and *FGFR2*. **(C)** TGF-β pathway genes. Note the expression of *ID2*, a marker of lung tip progenitors, and other genes known to be involved in lung development such as *BMP7, SMADs*, and *PPP2CA*. **(D)** WNT pathway genes. Note expression of multiple *FZD* family genes as well as *WNT7B*, a key regulator of lung development. Combined, these data shows that the LGR5+ epithelial cells have both receptor (*FGFR1, FGFR2*), and ligand (*SHH, WNT7*, and others) activity. AE: average expression; PE: percent expression.

**Supplement Figure S5. SHH, FGF, TGF-β and WNT pathway genes expressed by the different mesenchymal clusters at GD80.** (A) SHH pathway genes. Note expression of targets of SHH such as *SMO, GLI1, GLI2, PTCH1* and *PTCH2* among others (B) FGF pathway genes TGF-β pathway genes. Note expression of both *FGF10* as well as *FGFR1* and *SPRY2*, all key genes in lung development. This pattern of expression is consistent with that of immature1 cluster fetal lung mesenchymal cells (56). (C) TGF-β pathway genes. Note expression of *ID2* a marker of lung tip progenitors, and other genes known to be involved in lung development such as *BMP4, BMP7, SMADs*, and *PPP2CA*. (D) WNT pathway genes. Note expression of *LGR4, LGR5*, multiple *PRICKLE* genes as well as *SERPINF1* key regulator of lung development. Combined, these data shows that the LGR5^+^ mesenchymal cells have both receptor (*SMO, FGFR1, LGR4*, and *LGR5*) and ligand functions (*FGF10, TGFB1, TGFB2, WNT5A, WNT11* and others). AE: average expression; PE: percent expression.

**Supplement Figure S6. Comparison of transcriptome of organoid-derived LGR5^+^ and LGR5^-^ cells and analysis of markers for nerve associated fibroblasts and Schwan cell precursors.** (A) Organoid-derived LGR5^+^ and LGR5^-^ cells were sorted as described in materials and methods and subjected to bulk RNA sequencing analysis. Differential gene expression of key markers of airway epithelial cells are shown. LGR5^+^ cells expressed markers of both bud tips (*SFTPC*) and basal cells (*KRT14*), in addition to upregulating *AXIN2, BMP4* and *TGFB2*. (B1-B2) Volcano plot for differentially expressed genes in LGR5^+^ versus LGR5^-^ lung mesenchymal cells that are known markers of nerve-associated fibroblasts (perineurial, epineurial, and endoneurial), or Schwann cells (Schwann cells, non-myelinating SC, and myelinating SC). Markers used to differentiate the two populations are listed in Supplement Table 3. Genes in Table S3 missing from the volcano plots were either not expressed or not differentially expressed.

**Supplement Figure S7. Mapping of key pig and human LGR5^+^ signaling pathways to the adult human lung.** We have shown that both pigs and human postnatal LGR5^+^ mesenchymal cells have active SHH, TGF-β, FGF and WNT signaling pathways. To determine whether this signaling signature is unique to LGR5^+^/peribronchial fibroblasts we developed a pathway score composed of the genes that were differentially upregulated in both human and pigs (concordant genes only) and mapped those genes subsets to the Madissoon et al (10) scRNA-seq generated human atlas. (A). LGR5^+^ peribronchial fibroblasts (black oval) are located adjacent to myofibroblasts (purple arrow). Other mesenchymal clusters include smooth muscle cells (green arrow), an a broad fibroblasts cluster that includes perichondrial fibroblasts, alveolar fibroblast, adventitial fibroblast, and immune recruiting fibroblast (yellow arrow) as well as a cycling fibroblast cluster (pink arrow) (B) SHH pathway score was generated with genes *BOC, DZIP1, EVC2, GLI3, GPC6, HHAT, HHIP, KIF3A, PTCH1, SMO, SMURF2* and *SUFU* (C) TGF-β pathway score was generated with genes *ACVR2A, BMP5, BMP7, BMPR1A, INHBA, LTBP1,SMAD5, SMAD9* and *ZFYVE16*. (C) FGF signaling score was generated with *FGF2, FGF7, FGF10*, and *FGFR1* (E) Canonical WNT pathway score was generated with genes *AXIN2, CTNNB1, GSK3B, LEF1*, and *TCF4*. (F) Non-Canonical WNT/PCP (planar cell polarity) pathways score was generated with genes *DAAMF1, DAAM2, DVL1, DVL2, PRICKLE1, PRICKLE2, ROR1* and *ROR2*. In all cases, except for the canonical WNT pathway (F), the LGR5^+^/peribronchial fibroblast cluster had the strongest signaling of all the mesenchymal clusters.

**Supplement Figure S8. Mapping of nerve-associated fibroblasts markers to pig GD80, and human fetal and adult lung.** We have shown that LGR5 mesenchymal cells are spatially organized near nerve fibers surrounding the airways. We also showed that they have transcriptional signatures enriched for nerve-associated fibroblasts (NAF; Supplement Figure 5B). Using these data, we developed a NAF score using genes *OSR2, BMP4, FMOD, A2M, FN1, DPT, CRISPLD2, C3, AEBP1, BMP7, ETV1, PI16, APOD, ADAMTS5*, and *USP54* and mapped them jointly to multiple scRNA-seq atlases including our own GD80 dataset. (A1-A2). At GD80, the adult NAF score mapped to multiple cell clusters but was enriched in Mes 6, Mes 3 and Mes 4 (arrows). In contrast Mes 7 and the Epi cluster had minimal signal. (B1-B3) Analysis of the human fetal lung cell atlas dataset generated using He et al (65) showed that it includes LGR5 cells in epithelial clusters 3 and 5, as well as the fibroblasts cluster 0, identified by arrows and cluster numbers. (C1- C3) Mapping of NAF score to adult human atlas generated using Madissoon et al (10) dataset identified the LGR5+ peribronchial population (C1, arrow) overlapping with LGR5^+^ cells (C2, arrow) and NAF score populations (C3, arrow).

**Supplement Table S1.** Gene Set Enrichment Analysis for cell type (Module C8), Gene Ontology:Biological Process, and Gene Ontology:Molecular function using the top 500 upregulated genes in LGR5 PD29-38 mesenchymal cells.

**Supplement Table S2.** Summary of markers of nerve-associated cells and Schwann cells used to generate Supplement Figure 2.

**Supplement Table S3.** Expression in PD29-38 pig LGR5+ versus LGR5- mesenchymal cells of markers of distal human airway LGR5 mesenchymal cells identified by Murthy et al (11)

**Supplement Data Table S1.** Top 500 Upregulated genes in postnatal (PD29-38) LGR5^+^ mesenchymal cells.

**Supplement Data Table S2.** scRNAseq_GD80_AllMarkers

**Supplement Video S1.** Fetal and adult lung light-sheet microscopy.

