## Supplementary figures and images for "Novel porcine model reveals two distinct LGR5 cell types during lung development and homeostasis"

### Figure S1

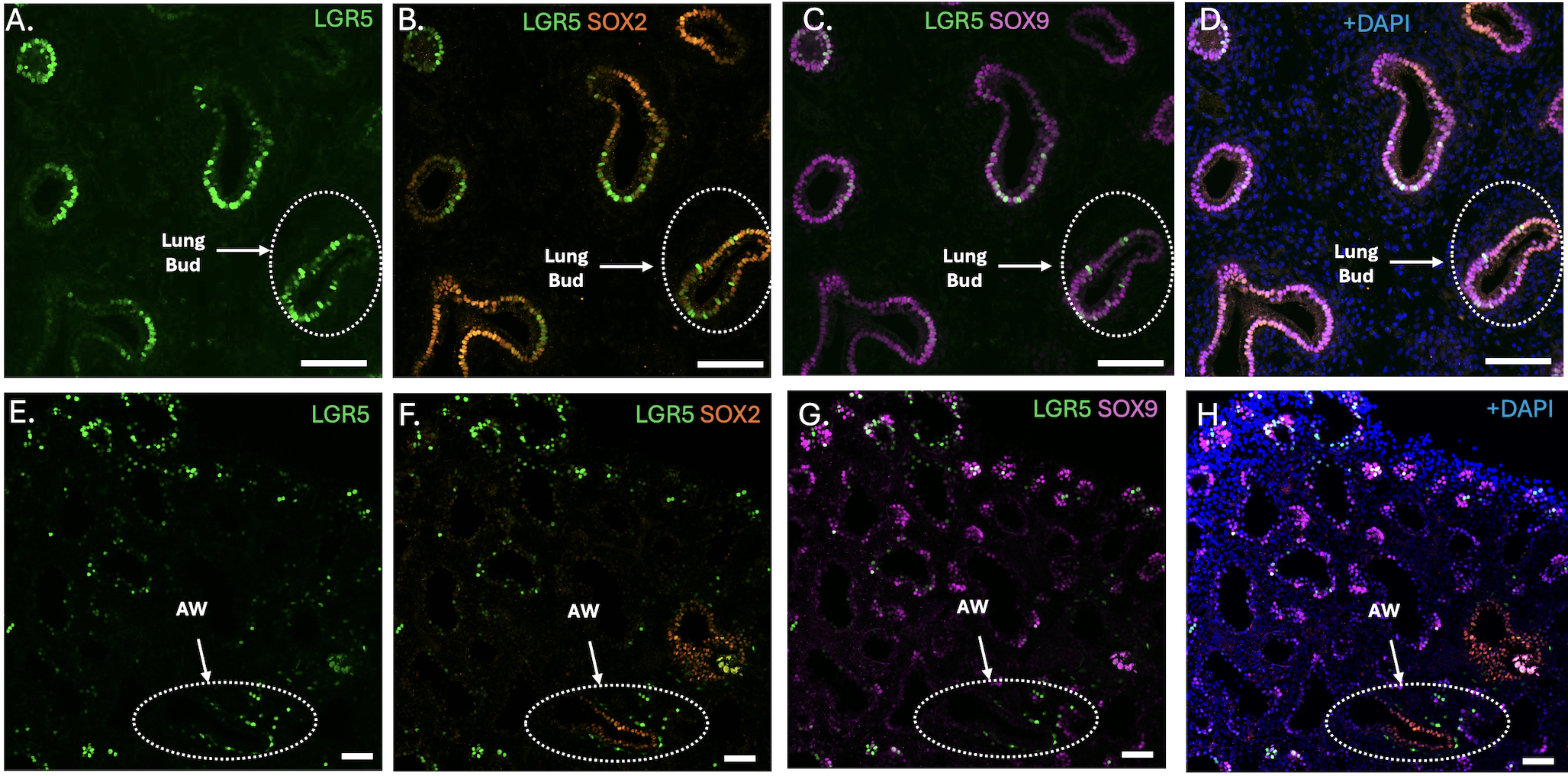

### Figure S2

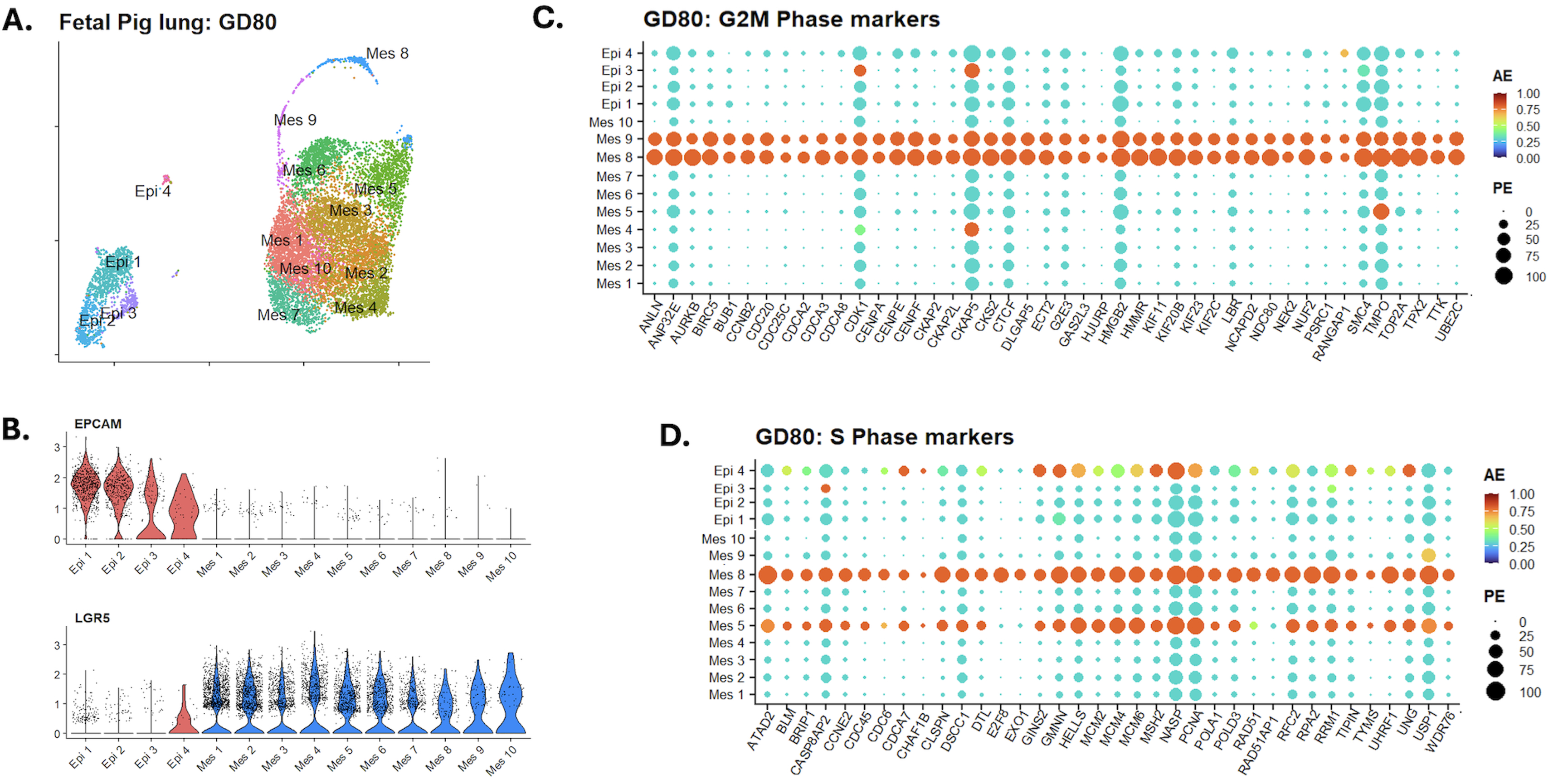

### Figure S3

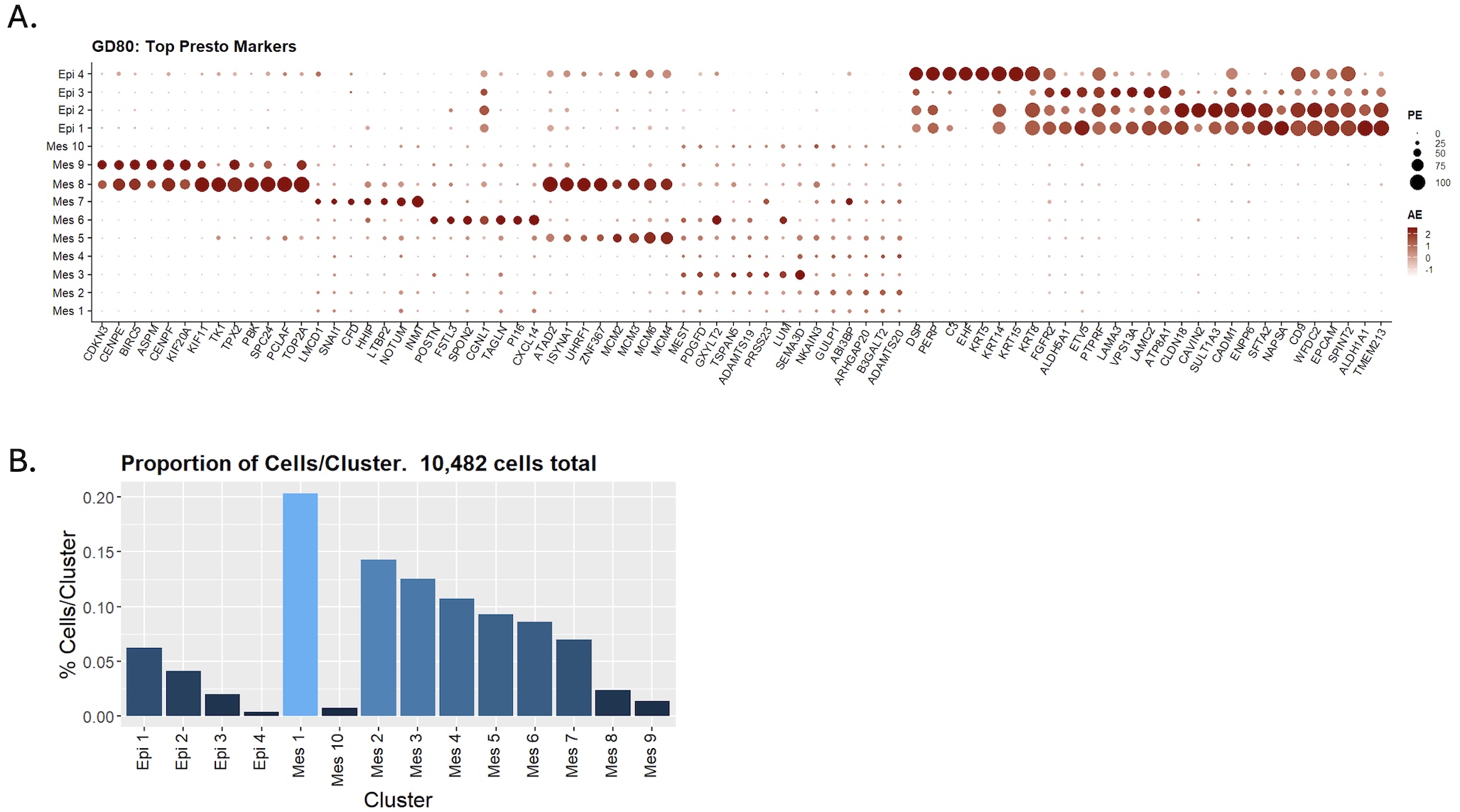

### Figure S4

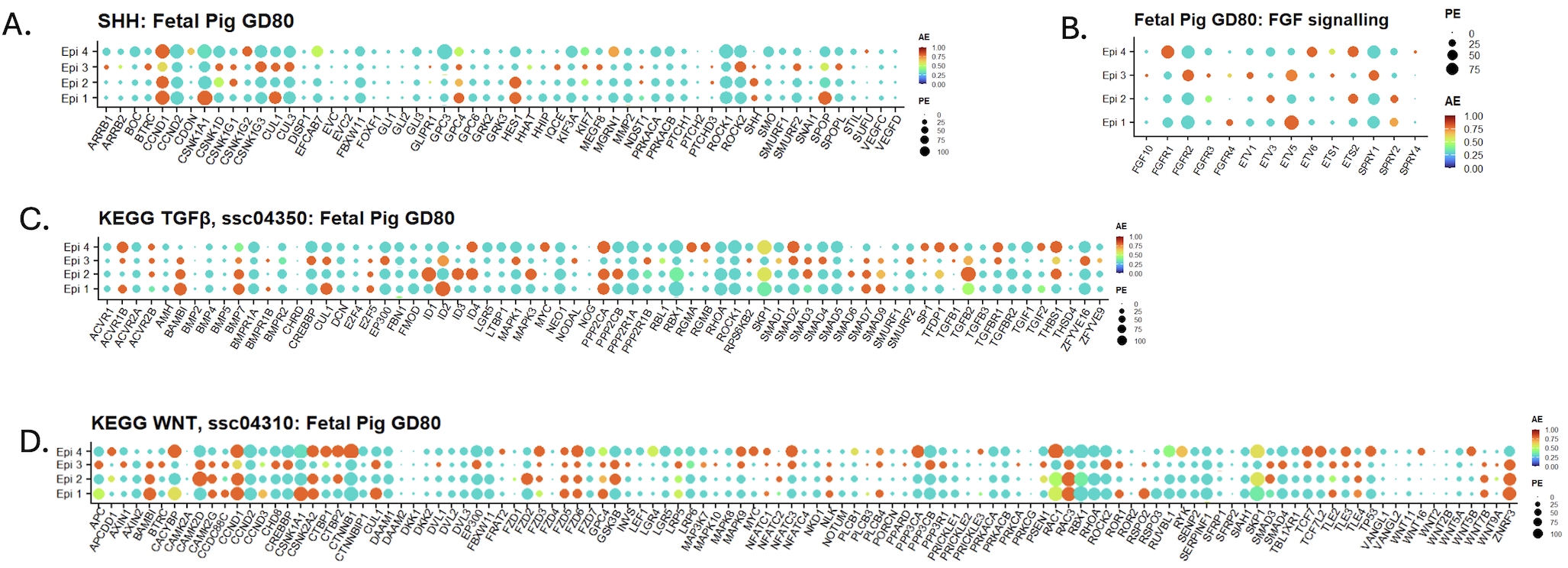

### Figure S5

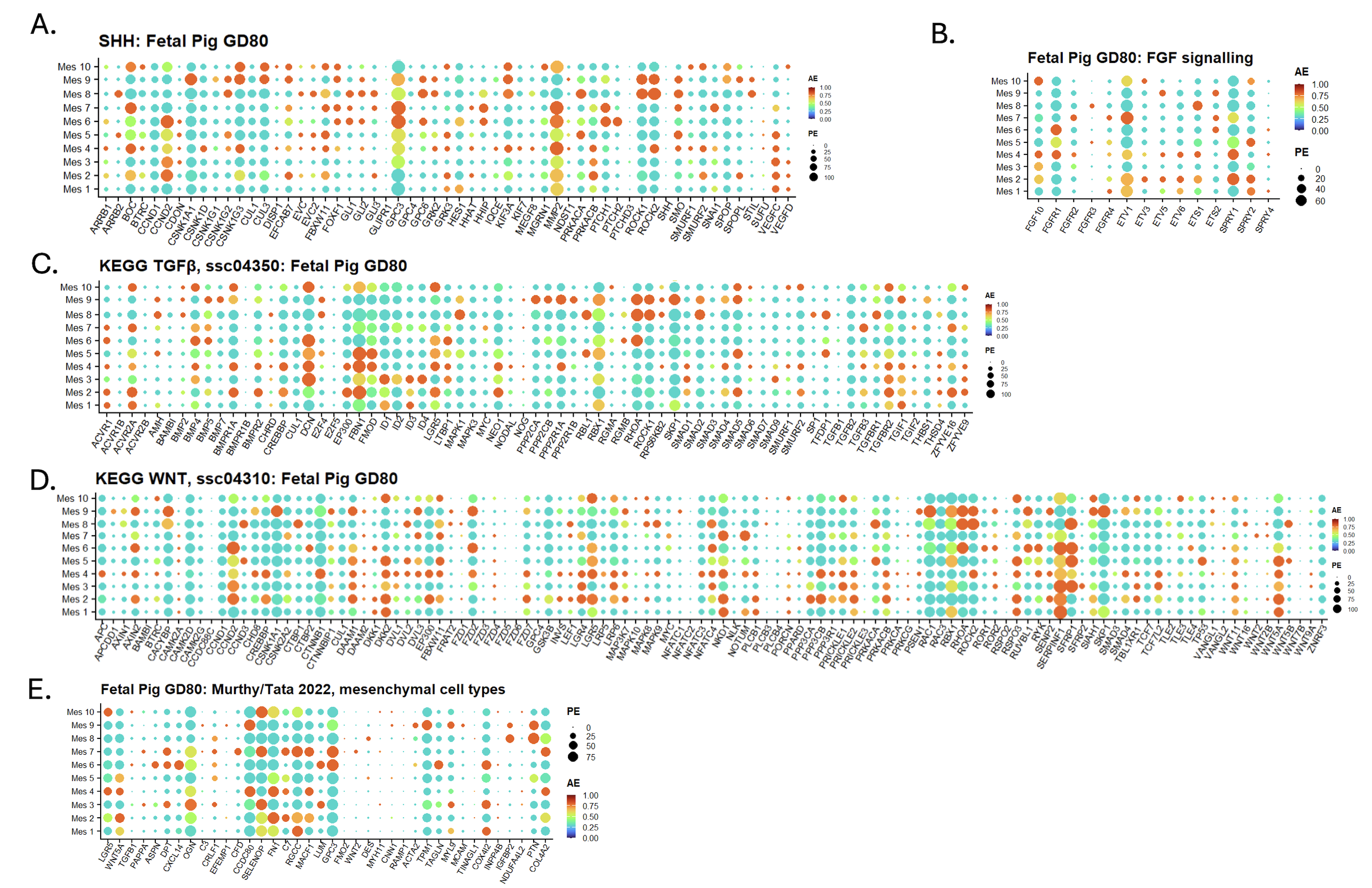

### Figure S6

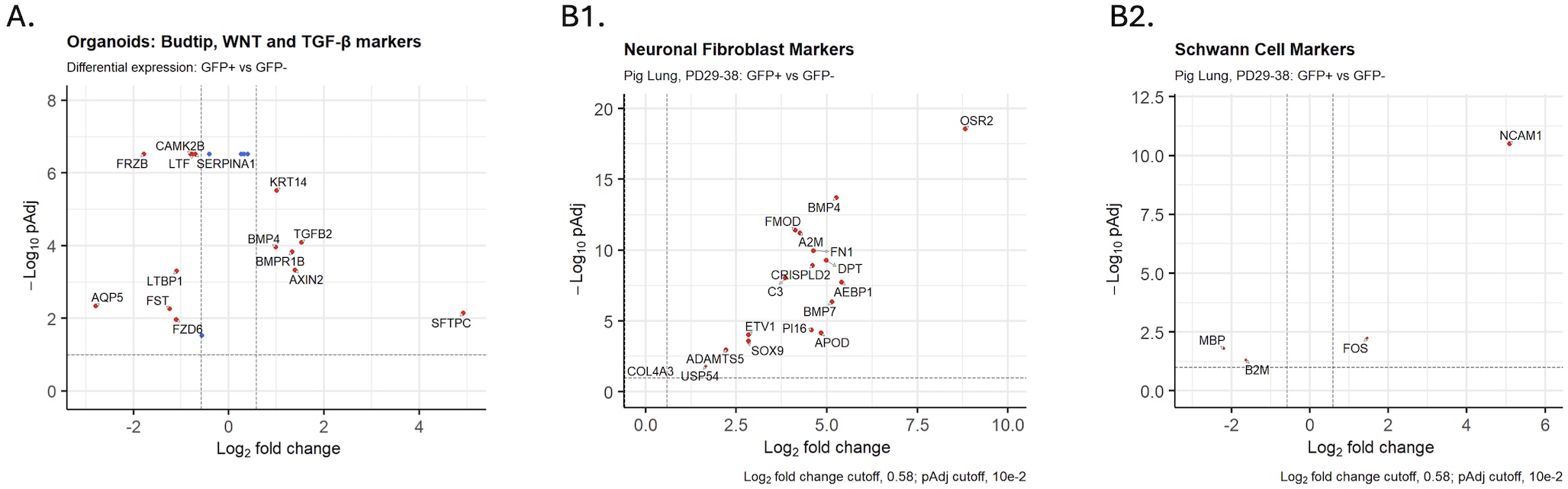

### Figure S7

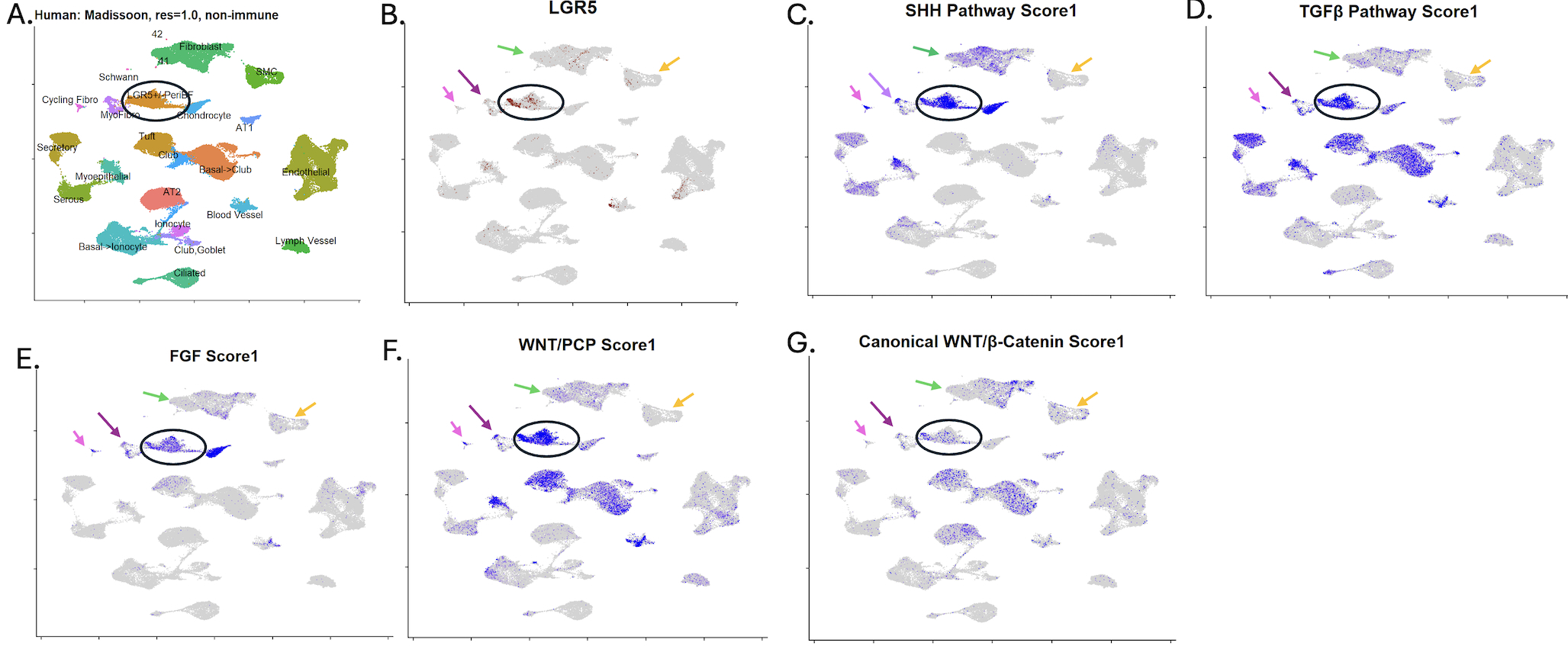

### Figure S8

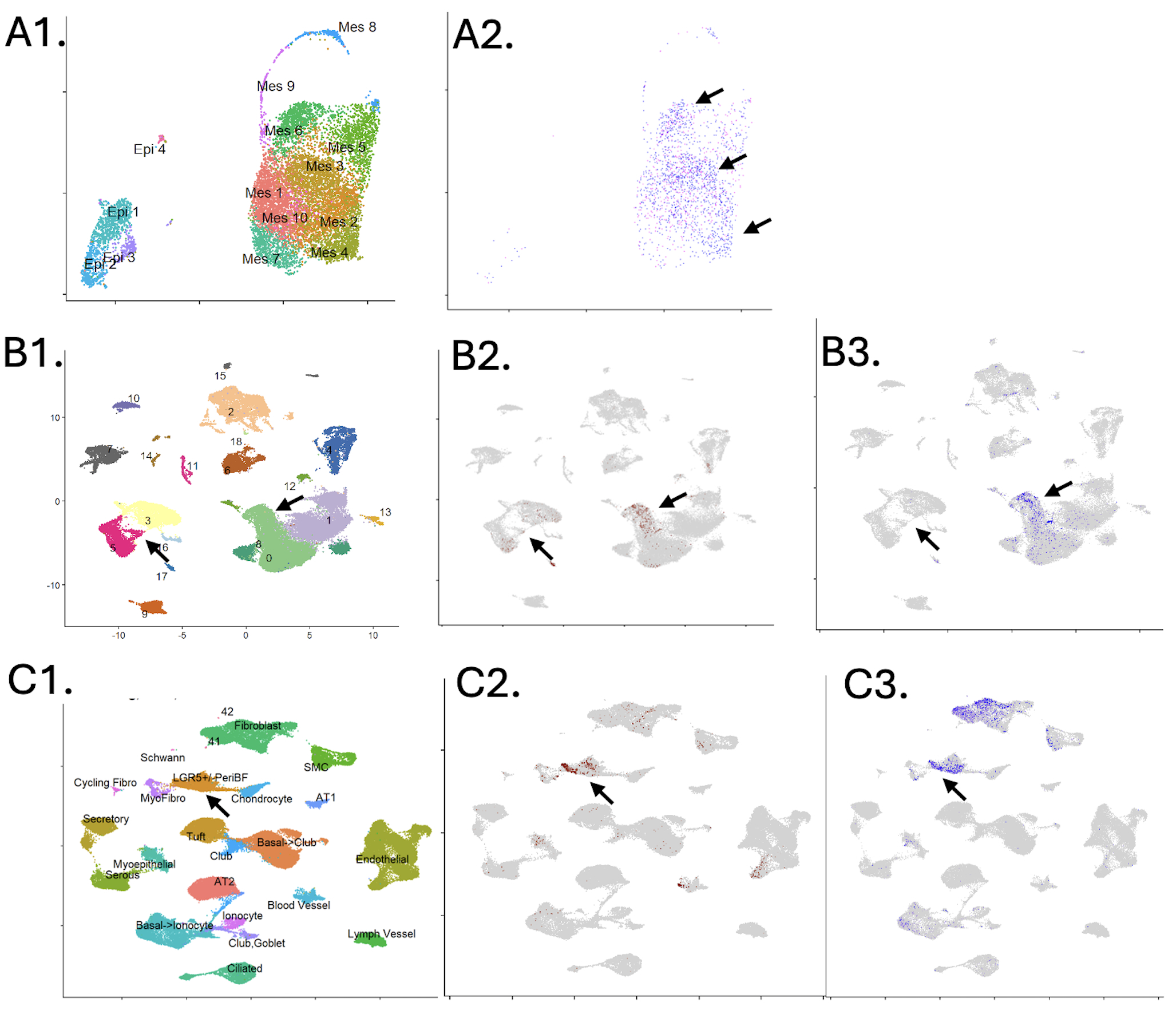
