## Supplementary material for "Novel porcine model reveals two distinct LGR5 cell types during lung development and homeostasis": Table S1

**Supplement Table S1. Gene Ontology and Cell type: Genes Upregulated in PD29-38 LGR5+ vs LGR5- Lung Mesenchymal cells.**

**A. GSEA: Cell Type**

Overlaps shown: 20

Genesets in collections: 830

Genes in comparison (n): 500

Genes in universe (N): 42,722

| Gene Set Name | k/K | p-value | FDR q-value |
|---|---|---|---|
| HAY_BONE_MARROW_STROMAL | 0.2243 | 1.49E-173 | 1.24E-170 |
| GAO_LARGE_INTESTINE_ADULT_CJ_IMMUNE_CELLS | 0.2495 | 3.70E-131 | 1.54E-128 |
| MURARO_PANCREAS_MESENCHYMAL_STROMAL_CELL | 0.1674 | 5.76E-97 | 1.59E-94 |
| CUI_DEVELOPING_HEART_C3_FIBROBLAST_LIKE_CELL | 0.5556 | 1.32E-94 | 2.73E-92 |
| DESCARTES_FETAL_THYMUS_STROMAL_CELLS | 0.4818 | 2.02E-90 | 3.34E-88 |
| HU_FETAL_RETINA_FIBROBLAST | 0.2234 | 6.04E-84 | 8.36E-82 |
| TRAVAGLINI_LUNG_ADVENTITIAL_FIBROBLAST_CELL | 0.2669 | 9.37E-84 | 1.11E-81 |
| DESCARTES_FETAL_EYE_STROMAL_CELLS | 0.5495 | 1.84E-72 | 1.91E-70 |
| AIZARANI_LIVER_C21_STELLATE_CELLS_1 | 0.3196 | 3.43E-71 | 3.16E-69 |
| GAO_LARGE_INTESTINE_24W_C1_DCLK1POS_PROGENITOR | 0.4848 | 5.40E-66 | 4.49E-64 |
| RUBENSTEIN_SKELETAL_MUSCLE_FAP_CELLS | 0.3184 | 2.34E-65 | 1.77E-63 |
| TRAVAGLINI_LUNG_ALVEOLAR_FIBROBLAST_CELL | 0.3354 | 1.58E-64 | 1.09E-62 |
| FAN_EMBRYONIC_CTX_BRAIN_ENDOTHELIAL_2 | 0.2145 | 4.84E-62 | 3.09E-60 |
| MANNO_MIDBRAIN_NEUROTYPES_HPERIC | 0.1145 | 1.01E-61 | 5.96E-60 |

| | | | |
|---|---|---|---|
| DESCARTES_FETAL_ADRENAL_STROMAL_CELLS | 0.3377 | 3.26E-61 | 1.81E-59 |
| DESCARTES_FETAL_KIDNEY_STROMAL_CELLS | 0.3425 | 3.05E-59 | 1.58E-57 |
| RUBENSTEIN_SKELETAL_MUSCLE_FBN1_FAP_CELLS | 0.2111 | 9.17E-58 | 4.48E-56 |
| DURANTE_ADULT_OLFACTORY_NEUROEPITHELIUM_FIBROBLASTS_STROMAL_CELLS | 0.4878 | 2.95E-55 | 1.36E-53 |
| FAN_OVARY_CL6_PUTATIVE_EARLY_ATRETIC_FOLLICLE_THECAL_CELL_2 | 0.1933 | 1.40E-52 | 6.11E-51 |
| HE_LIM_SUN_FETAL_LUNG_C3_OMD_POS_ENDOTHELIAL_CELL | 0.1577 | 1.21E-45 | 5.03E-44 |

**B. GSEA - Gene Ontology: Biological Processes**

Overlaps shown: 20
Genesets in collections: 7,647
Genes in comparison (n): 500
Genes in universe (N): 42,722

| GENE SET NAME | k/K | p-value | FDR q-value |
|---|---|---|---|
| GOBP_ANIMAL_ORGAN_MORPHOGENESIS | 0.0998 | 7.53E-65 | 5.76E-61 |
| GOBP_CIRCULATORY_SYSTEM_DEVELOPMENT | 0.0842 | 5.46E-57 | 1.64E-53 |
| GOBP_TUBE_DEVELOPMENT | 0.0864 | 6.43E-57 | 1.64E-53 |
| GOBP_EXTERNAL_ENCAPSULATING_STRUCTURE_ORGANIZATION | 0.1815 | 8.68E-52 | 1.66E-48 |
| GOBP_CELL_ADHESION | 0.0699 | 2.17E-51 | 3.32E-48 |
| GOBP_TUBE_MORPHOGENESIS | 0.0881 | 9.05E-48 | 1.15E-44 |
| GOBP_RESPONSE_TO_ENDOGENOUS_STIMULUS | 0.0608 | 4.49E-46 | 4.91E-43 |
| GOBP_VASCULATURE_DEVELOPMENT | 0.0925 | 2.93E-45 | 2.80E-42 |
| GOBP_RESPONSE_TO_GROWTH_FACTOR | 0.0969 | 7.33E-45 | 6.23E-42 |
| GOBP_ANATOMICAL_STRUCTURE_FORMATION_INVOLVED_IN_MORPHOGENESIS | 0.0715 | 9.59E-44 | 7.33E-41 |
| GOBP_ENZYME_LINKED_RECEPTOR_PROTEIN_SIGNALING_PATHWAY | 0.0802 | 1.64E-43 | 1.14E-40 |
| GOBP_CELL_MOTILITY | 0.0579 | 8.03E-43 | 5.11E-40 |
| GOBP_SKELETAL_SYSTEM_DEVELOPMENT | 0.1153 | 2.05E-41 | 1.21E-38 |
| GOBP_NEUROGENESIS | 0.0583 | 5.95E-41 | 3.25E-38 |
| GOBP_CELL_CELL_SIGNALING | 0.056 | 7.78E-38 | 3.97E-35 |
| GOBP_EMBRYO_DEVELOPMENT | 0.0701 | 1.06E-37 | 5.06E-35 |
| GOBP_REGULATION_OF_CELL_DIFFERENTIATION | 0.0571 | 2.81E-37 | 1.26E-34 |
| GOBP_GENERATION_OF_NEURONS | 0.0586 | 1.24E-35 | 5.27E-33 |
| GOBP_NEGATIVE_REGULATION_OF_RESPONSE_TO_STIMULUS | 0.0536 | 1.88E-35 | 7.56E-33 |
| GOBP_BLOOD_VESSEL_MORPHOGENESIS | 0.0874 | 5.21E-35 | 1.90E-32 |

**C. GSEA - Gene Ontology: Molecular Function**

Overlaps shown: 20
Genesets in collections: 1,799
Genes in comparison (n): 500
Genes in universe (N): 42,722

| GENE SET NAME | k/K | p-value | FDR q-value |
|---|---|---|---|
| GOMF_EXTRACELLULAR_MATRIX_STRUCTURAL_CONSTITUENT | 0.3193 | 7.26E-61 | 1.31E-57 |
| GOMF_GLYCOSAMINOGLYCAN_BINDING | 0.1818 | 8.28E-39 | 7.45E-36 |
| GOMF_SIGNALING_RECEPTOR_BINDING | 0.0589 | 3.31E-36 | 1.98E-33 |
| GOMF_HEPARIN_BINDING | 0.2069 | 4.62E-34 | 2.08E-31 |
| GOMF_SULFUR_COMPOUND_BINDING | 0.148 | 1.81E-32 | 6.51E-30 |
| GOMF_STRUCTURAL_MOLECULE_ACTIVITY | 0.0754 | 7.39E-31 | 2.22E-28 |
| GOMF_INTEGRIN_BINDING | 0.1688 | 1.56E-22 | 4.01E-20 |
| GOMF_COLLAGEN_BINDING | 0.2794 | 2.79E-21 | 6.28E-19 |
| GOMF_EXTRACELLULAR_MATRIX_STRUCTURAL_CONSTITUENT_CONFERRING_TENSILE_STRENGTH | 0.3556 | 4.62E-20 | 9.23E-18 |
| GOMF_PROTEIN_CONTAINING_COMPLEX_BINDING | 0.0403 | 1.18E-19 | 2.12E-17 |
| GOMF_CELL_ADHESION_MOLECULE_BINDING | 0.0663 | 3.07E-17 | 5.01E-15 |
| GOMF_GROWTH_FACTOR_BINDING | 0.1471 | 2.02E-16 | 3.03E-14 |
| GOMF_MOLECULAR_TRANSDUCER_ACTIVITY | 0.0386 | 1.38E-15 | 1.91E-13 |
| GOMF_CALCIUM_ION_BINDING | 0.0536 | 1.09E-14 | 1.41E-12 |
| GOMF_SIGNALING_RECEPTOR_REGULATOR_ACTIVITY | 0.0585 | 1.40E-13 | 1.68E-11 |
| GOMF_TRANSMEMBRANE_RECEPTOR_PROTEIN_KINASE_ACTIVITY | 0.1728 | 6.73E-13 | 7.57E-11 |
| GOMF_WNT_PROTEIN_BINDING | 0.3125 | 2.26E-12 | 2.39E-10 |
| GOMF_PEPTIDASE_REGULATOR_ACTIVITY | 0.0881 | 3.93E-12 | 3.92E-10 |
| GOMF_PLATELET_DERIVED_GROWTH_FACTOR_BINDING | 0.6364 | 9.14E-12 | 8.66E-10 |
| GOMF_GROWTH_FACTOR_ACTIVITY | 0.1049 | 1.00E-11 | 9.01E-10 |
