## Supplementary material for "Novel porcine model reveals two distinct LGR5 cell types during lung development and homeostasis": Table S2

### Supplement Table S2. Markers of nerve and nerve-associated cells

| Marker | Location | Cell Type | Reference |
|---|---|---|---|
| A2M | PNS | EnC | 44 |
| ABLIM1 | CNS | NSC | 52 |
| ADAMTS5 | PNS | EpN | 44 |
| AEBP1 | PNS | PNAF | 46 |
| ANGPT1 | CNS | NSC | 52 |
| ANTXR1 | CNS | NSC | 52 |
| APOD | PNS | PNAF | 46 |
| APOE | PNS | mySC | 46,47 |
| BGN | PNS | MNC | 48 |
| BMP4 | PNS | EpN | 4,48 |
| BMP7 | PNS | EnC | 44 |
| C3 | PNS | PNAF | 10,46 |
| CARMIL1 | CNS | NSC | 52 |
| CHPT1 | CNS | NSC | 52 |
| CLEC3B | PNS | PNAF | 46 |
| CLU | CNS | NSC | 52 |
| COL11A1 | PNS | EnC | 44 |
| COL4A3 | PNS | PnC | 44 |
| COL5A1 | CNS | NSC | 52 |
| CPE | CNS | VLMC | 52 |
| CRISPLD2 | PNS | EpN | 44 |
| CTNND2 | CNS | NSC | 52 |
| DAPK1 | CNS | VLMC | 52 |
| DPT | PNS | EpN, PNAF | 10,44,46 |
| EFEMP1 | CNS | NSC | 52 |
| ETV1 | PNS | EnP | 44 |
| F3 | CNS | NSC | 51 |
| FABP7 | PNS | NP | 49 |
| FBLN7 | PNS | EpN | 44 |
| FMOD | PNS | EpN | 44,48 |
| FN1 | PNS | PNAF | 46 |
| FOS | PNS | mySC | 46 |
| GSN | PNS | PNAF | 46 |

| | | | |
|---|---|---|---|
| ITGA6 | PNS | PnC | 44,50 |
| LGALS3 | PNS | MNC | 48 |
| LMO7 | PNS | PnC | 44 |
| LTBP1 | CNS | NSC, VLMC | 52 |
| MBP | PNS | mySC | 44,46 |
| MFGE8 | CNS | NSC | 52 |
| NCAM1 | PNS | nmSC | 44 |
| NCMAP | PNS | mySC | 44,50 |
| NDRG2 | CNS | NSC | 52 |
| NEBL | CNS | NSC, ImN, OPC | 52 |
| NES | PNS | NP, mySC | 44,49 |
| NGFR | PNS | NSC, nmSC, mSC | 44,46,48 |
| NHSL1 | CNS | NSC | 52 |
| NTRK2 | CNS | NSC, ExN, ImN, OPC | 52 |
| OSR2 | PNS | EnC, EnP, nmSC | 44,46 |
| PALM | CNS | NSC, OL | 52 |
| PI16 | PNS | PNAF | 46 |
| PRX | PNS | mSC | 50 |
| PTPN13 | CNS | NSC, VLMC | 52 |
| REV3L | CNS | NSC, OPC | 52 |
| RORA | CNS | NSC, VLMC | 52 |
| SDC2 | CNS | NSC | 52 |
| SORBS1 | PNS | EpN | 44,50 |
| SOX5 | PNS | EnP | 44 |
| SOX6 | PNS | EnP | 44 |
| SOX8 | PNS | EnP | 44 |
| SOX9 | PNS | EnP | 44 |
| STXBP6 | PNS | EpN | 50 |
| TGFB2 | CNS | NSC | 52 |
| THY1 | PNS | EnC | 44 |
| TIAM1 | PNS | PNAF | 50 |
| TPCN1 | CNS | NSC | 52 |
| USP54 | PNS | PNAF, EnC | 44,50 |
| VAT1L | CNS | NSC | 52 |
| WWC1 | CNS | NSC | 52 |
| ZFHX4 | CNS | NSC | 52 |

| | | | |
|---|---|---|---|
| ZNRF3 | CNS | NSC | 52 |

EnC = Endoneurial cell
EnP = Endoneurial precursor
EpC = Epineurial cell
ExN = Excitatory neuron
ImN = Immature neuron
NP = Neural progenitor
nmSC = Non-myelinating Schwann cell
mSC = Myelinating Schwann cell
NSC = Neural stem cell
OL = Oligodendrocyte
OPC = Oligodendrocyte progenitor cell
PnC = Perineurial cells
PNAF = Peripheral nerve associated fibroblast
SC = Schwann cell
VLMC = Vascular leptomeningeal cell
