## Supplementary material for "Novel porcine model reveals two distinct LGR5 cell types during lung development and homeostasis": Table S3

Supplement Table S3. Expression in PD29-38 pig LGR5+ versus LGR5- mesenchymal cells of markers of distal human airway LGR5 mesenchymal cells identified by Murthy et al (11)

| Gene | baseMean | log2FoldChange | padj | SumPos | SumNeg | DiffCnt |
|---|---|---|---|---|---|---|
| WNT5A | 299.8774261 | 4.052966508 | 4.85E-10 | 2622 | 172 | 2450 |
| TGFBI | 4279.792069 | 4.057273738 | 5.38E-11 | 37373 | 2017 | 35356 |
| PAPPA | 216.9449047 | 2.322003639 | 0.074846166 | 1595 | 232 | 1363 |
| ASPN | 2665.48191 | 6.428827904 | 1.99E-25 | 24269 | 301 | 23968 |
| DPT | 565.74937 | 4.9851637 | 5.36E-10 | 5107 | 170 | 4937 |
| CXCL14 | 338.3330372 | 5.111290869 | 1.67E-15 | 3096 | 87 | 3009 |
| OGN | 4582.619929 | 4.862577796 | 4.84E-10 | 41022 | 1620 | 39402 |
| C3 | 7172.829303 | 3.856332923 | 9.41E-09 | 64026 | 4589 | 59437 |
| WNT2 | 24.30290223 | 2.613790429 | 0.076143531 | 198 | 28 | 170 |
| DES | 17.39296307 | 0.916257237 | 0.682603542 | 87 | 60 | 27 |
| MYH11 | 131.2729789 | -1.966884895 | 0.000673335 | 210 | 941 | -731 |
| CNN1 | 6.854517778 | -1.473523998 | 0 | 14 | 39 | -25 |
| RAMP1 | 37.44509067 | -2.062907452 | 0.048259891 | 61 | 277 | -216 |

SumPos refers to LGR5+ cells and SumNeg to LGR5- cells.
